# Localized Reconstruction of Multimodal Distance Distribution from DEER Data of Biopolymers

**DOI:** 10.1101/2025.01.02.631084

**Authors:** Karen Tsay, Timothy Keller, Yann Fichou, Jack H. Freed, Song-I Han, Madhur Srivastava

## Abstract

Pulsed Dipolar ESR Spectroscopy (PDS) is a uniquely powerful technique to characterize the structural property of intrinsically disordered proteins (IDPs) and polymers and the conformational evolution of IDPs and polymers, e.g. during assembly, by offering the probability distribution of segment end-to-end distances. However, it is challenging to determine distance distribution *P*(*r*) of IDPs by PDS because of the uncertain and broad shape information that is intrinsic to the distance distribution of IDPs. We demonstrate here that the Srivastava-Freed Singular Value Decomposition (SF-SVD) point-wise mathematical inversion method along with wavelet denoising (WavPDS) can aid in obtaining reliable shapes for the distance distribution, *P*(*r*), for IDPs. We show that broad regions of *P*(*r*) as well as mixed narrow and broad features within the captured distance distribution range can be effectively resolved and differentiated without *a priori* knowledge. The advantage of SF-SVD and WavPDS is that the methods are transparent, requiring no adjustable parameters, the processing of the magnitude for the probability distribution is performed separately for each distance increment, and the outcome of the analysis is independent of the user’s judgement. We demonstrate the performance and present the application of WavPDS and SF-SVD on model ruler molecules, model polyethylene glycol polymers with end-to-end spin labeling, and IDPs with pairwise labeling spanning different segments of the protein tau to generate the transparent solutions to the *P*(*r*)’s including their uncertainties and error analysis.

## 1. INTRODUCTION

Pulsed Dipolar Electron Spin Resonance Spectroscopy (PDS) is a major tool in structural biology and has seen substantial growth in recent years for structural studies of biomacromolecules and their complexes.^1–14^ The strength of PDS is that it can be utilized to obtain structural information of macromolecules of virtually any size and complexity, and independent of whether the part to be resolved is structured or intrinsically disordered, as long as a pair of spin labels (commonly nitroxide-based) can be attached at two specified sites (including the two ends) without significantly perturbing the structural property of the macromolecule. The major and unique strength of PDS is its ability to obtain not only the mean distance, *r*, between the pair of spin labels on the macromolecule, but also the probability distribution of distances, *P*(*r*).^1,2,8,15–19^ This renders PDS an effective characterization tool of disordered and dynamical proteins with multiple conformations, as the shape of *P*(*r*) offers information about whether a defined distance, multiple distances or a broad ensemble of distances are populated. However, the full potential of PDS for obtaining *P*(*r*) remains underutilized, as most analysis methods for PDS are to derive single or multiple discrete mean distances from time-domain data acquired by PDS. It is a major challenge to reliably determine the shape of the distance distribution *P*(*r*) with quantifiable uncertainty, especially if the number, width and shape of the different populations are not known *a priori*. Hence, it is essential to use unbiased and transparent distance reconstruction methods, in which the determination of the key results and their uncertainties for the *P*(*r*) is inherent to the method and transparent, and hence independent of user-adjustable parameters or competence.

In PDS, a dipolar evolution function, *S*(*t*), is acquired that is modulated by the interaction between an ensemble of spin label pairs. The distance distribution, *P*(*r*), can be obtained from *S*(*t*) with knowledge of the kernel, *K*, that is the weighing function for the dipolar coupling as a function of *t* and *r*, averaged over θ between the inter-spin dipolar vector and *B*_0_. The mathematical form is represented as *KP* = *S*, where *S* is the dipolar evolution function that is contained in *V*(*t*), the experimental PDS signal, *K* is known and *P* needs to be obtained. Here, *P* is a function of *r*, hence *P*(*r*) and *K* is a function of *r* and time, *t*, hence *K*(*t, r*). However, the solution cannot be simply obtained in the form *P* = *K*^−1^*S* because this is a mathematically ill-posed problem, meaning that the calculation of *P* using that form yields an unstable solution that does not reliably represents the actual distance distribution. The resolution of the distance distribution is further limited by the presence of noise in *V*(*t*).

Furthermore, when *represents* a superposition of well-defined narrow functions, but is a broad distribution that represent intramolecular distances of intrinsically disordered proteins (IDPs) or polymers tagged with a pair of spin labels, then the ill-posed nature of the problem and sensitivity limitations are aggravated because a much greater ensemble of distances must be accurately captured, while the signal is also distributed across a much broader distance range, hence diluting the signal for a given distance increment. An analysis method of time-domain dipolar signal that effectively reconstructs the distance distribution of IDPs or polymers should meet the following criteria:

1. It can determine and differentiate distance distributions containing distinct populations with multiple distances and with both broad and narrow features, as well as a single broad distribution representing a dynamic ensemble.
2. It can obtain the distance distribution despite a substantial noise level in the experimental dipolar signal, *V*(*t*). Note that the *V*(*t*) of IDPs readily become much noisier when using even marginally longer evolution times.

Methods, such as *a priori* model-based Gaussian fitting ^20,21^, training-based neural networks^22,23^, and model-free Tikhonov regularization (TIKR)^24–27^ can be used to obtain distance distributions, but they have one or more limitations that make them less-suited for structure studies of IDPs.

Model fitting methods, such as GLAADvu^20,21^ require *a priori* information about the shape and nature of the distance distributions. For non-descriptive shapes such as those of intrinsically disordered systems that may adopt partially ordered structures, the model fitting methods may at best be limited to determining mean distances over all the features. A similar problem exists for the training-based DEERNet approach^22,23^, where complete and/or reliable simulated data for a conformational ensemble of IDPs is typically unavailable, and is extremely costly to obtain, or in the case of partially ordered and/or large IDPs, cannot be reliably obtained with current computational tools. In other words, obtaining a large enough library to capture the entire conformational space of IDPs is illusive, especially as one does not know *a priori* whether the polymers or the IDPs of interest fit into well-defined polymer physics models, adopt distinct conformations because the polymer sequence is designed to partially fold, populate transient secondary structures and/or adopt partially ordered structures. TIKR^24–27^ has no *a priori* information or training requirement, but the single regularization parameter and its selection criteria limits the shape and resolution of the distance distribution. Reliable analysis using TIKR becomes especially problematic: 1) when both narrow and broad features are present in *P*(*r*), which cannot be simultaneously and optimally captured in a single *P*(*r*) because a single regularization parameter, *λ*, must be chosen that may only be optimal for one of the features, yielding a compromise between all the features present; and/or 2) when a distribution encompasses multiple features, but just appears as one broad distribution from using a too high *λ* value to deal with the presence of noise. With TIKR, the broad distribution can either reflect the true distribution of an ensemble of interconverting conformers or the smoothening out of a distribution that has narrower features (obscured by the presence of noise), both of which may rely on the use of the same regularization parameter. In any of these scenarios in which the shape of *P*(*r*) is more complex than a single narrow distribution, the solution for *P*(*r*) found by TIKR (among several candidate *P*(*r*)s) that best matches the experimental PDS signal depends on the choice for the regularization parameter that, in turn, can be neither rigorously nor transparently made, but rather depends on the intuition and experience of the user. For the determination of the shape of *P*(*r*) of IDPs, it is essential to have access to a method that offers rational and transparent criteria to differentiate between a broad distribution and one that contains single or multiple populations with mixed disorder.

We present that the Srivastava Freed-Singular Value Decomposition (SF-SVD) method is uniquely well suited to reconstruct the distance distribution of IDPs and polymers in a transparent and user-independent approach, where point-wise reconstruction of distance distribution is carried out so that the probability density obtained for each distance is independent of the analysis of the probability density of the other distances.^28,29^ The SF-SVD method obviates the need of a single regularization parameter as used in TIKR, but to use TIKR terminology, uses “regularization parameters customized for each distance increment or range”. In fact, it is actually free of any regularization parameters, but rather solves for the *convergent* value of *P*(*r*_*j*_) for each *j*, i.e. for each distance increment in real space. This enables the independent reconstruction of narrow and broad features of the distance distribution, without compromising the fidelity of one or the other. No regularization is needed; instead, convergent solutions are obtained. This approach does not use any criteria based on least square minimization, as is used in most regularization methods, including TIKR. This distance increment-by-increment reconstruction method of *P*(*r*) enables the generation of multimodal distance distributions containing features of varying width and shape, with resolution only limited by the number of *P*(*r*) entries that, in turn, is dictated by the number of data points constituting *S* and the duration of the echo decay in the time domain. The SF-SVD method ensures that a broad distribution (associated with the intrinsically disordered state) and multiple distributions within a broad range (associated with partial and dynamical structures of a heterogeneous ensemble of IDPs) can be distinguished. Of course, in the presence of noise, there is some uncertainty in the *P*(*r*_*j*_), which shows up as some fluctuations over its range of the *P*(*r*_*j*_) values obtained in the convergent region based on the number of singular value contributions (SVC’s). This defines *P*(*r*_*j*_) ‘s *uncertainty* and is independently obtained for each distance, thus yielding localized uncertainty for *P*(*r*_*j*_) for every given *r*_*j*_. For noisy data, the uncertainty becomes larger due to large variation in *P*(*r*_*j*_) values, even in the convergent region where *P*(*r*_*j*_) has converged as a function of SVCs.

Another important feature of SF-SVD is that it does not need to impose the constraint that *P*(*r*_*j*_) values be greater than 0, i.e. *P*(*r*_*j*_) > 0. The convergent *P*(*r*_*j*_) values always satisfy this condition. At the same time, negative values that originate from artifacts in experimental PDS signal, including those associated with data processing such as background correction and offset subtraction, are allowed and hence correctly reveal the effects of artifacts on the distance distribution without artificially truncating those fluctuations. For those cases where artifacts with negative values are substantial, data pre-processing and experiments need to be adjusted/repeated. For cases where artifacts are negligible, SF-SVD naturally generates a distance distribution with *P*(*r*_*j*_) > 0. In TIKR regularization, *P*(*r*) candidates are constrained to *P*(*r*) > 0, assuming that the PDS signal lacks artifacts, noise and/or other experimental effects. Often, there may not be any (or good) *P*(*r*) candidates that satisfy non-negativity constraints, in which case negative *P*(*r*) values need to be artificially truncated to 0. Commonly used pseudo SVD inversion methods to generate candidate *P*(*r*) solutions frequently encounter such problems.

As SF-SVD yields the *P*(*r*) from the dipolar evolution function, *S*, embedded in the experimental PDS signal, *V*(*t*), it is important to have *V*(*t*) with a high Signal-to-Noise Ratio (SNR). The problem of noise in the *V*(*t*) is overcome by applying the wavelet denoising method, WavPDS, which currently can recover signals for SNR ≥ 3 from random noise. The approach uses wavelet transforms to separate low magnitude wavelet coefficients representing noise from higher magnitude coefficients representing a coherent PDS time-domain signal. It has been used in several studies that demonstrate reliable analysis of double electron-electron resonance (DEER) data using WavPDS^4,6,30,31^. The use of wavelet denoising removes the random noise from PDS signal prior to applying SF-SVD to reconstruct *P*(*r*).

In this study, we test the utility of SF-SVD combined with WavPDS using DEER data of samples with a diversity of both known and unknown distance distributions, acquired with different collection and evolution times, with the goal to demonstrate that the shape of *P*(*r*) can be reliably determined. The segment end-to-end distance distribution, *P*(*r*), of a tau monomer in solution state and polymer end to end distance distribution, *P*(*r*), of polyethylene glycol (PEG), both show broad and smooth distributions, but their shapes are still distinctly different for the IDP and PEG. In contrast, the segment end-to-end distance distribution of fibrilized tau proteins adopts a distinctly non-smooth shape that contains both broad and narrow features, showcasing the benefit of SF-SVD with WavPDS in resolving localized distance distributions and enabling better baseline corrections.

## 2. METHOD

Details of the experimental setup and parameters, data processing workflow, theory of DEER data analysis, background correction, and the SF-SVD method for distance distribution reconstruction and uncertainty analysis are presented below.

### 2.1 Experimental Double Electron Electron Resonance (DEER)

The DEER experiments were performed with a pulsed Q-band Bruker E580 Elexsys spectrometer, equipped with a Bruker QT-II resonator and a 300 W TWT amplifier with an output power of 20 mW for the recorded data (Applied Systems Engineering, Model 177Ka). The temperature of the cavity was maintained at 65 K using a Bruker/ColdEdge FlexLine Cryostat (Model ER 4118HV-CF100). The bridge is equipped with an Arbitrary Waveform Generator to create shaped pulses for increased measurement fidelity and sensitivity. The samples were made in D_2_O buffer with 30 % (v/v) deuterated glycerol (used as the cryoprotectant)^18^. To perform an experiment, approximately 40 μL of sample is added to a 3 mm OD, 2 mm ID quartz capillary and flash frozen in liquid nitrogen to preserve the state of sample containing an ensemble of conformations by vitrification.

The following 4-pulse DEER sequence was applied to all samples: *π*_*obs*_/2 – *τ*_1_ – *π*_*obs*_ – (*t* − *π*_*pump*_) – (*τ*_2_ − *t*) – *π*_*obs*_ – *τ*_2_ – echo. *V*(*t*) is recorded as the integral of the refocused echo as a function of time delay, *t*, between the Hahn echo and pump pulse. Rectangular observe pulses and chirp pump pulse were used with the following pulse durations: *π*_*obs*_/2= 20 ns, *π*_*obs*_= 40 ns, *π*_*pump*_ = 100 ns. The chirp pump pulse was applied with a frequency width of 60 MHz to excite a distinct spin population, referred to as B spins, while the observe pulse was set 33 G up field from the center of the pump frequency range to probe another distinct spin population, the A spins. The *τ*_1_ value was set to 180 ns and *τ*_2_ was set according to the SNR of the dipolar signal. The data was acquired with a resolution of 16 or 32 ns, 16-step phase cycling, and signal averaging.

### 2.2 Data Processing Workflow

The workflow of distance reconstruction utilizing WavPDS and SF-SVD is given in Fig. 2. All the data presented in this paper were processed with the following procedures: (1) denoising with WavPDS^32^, (2) background correction with a logarithmic approach, and (3) reconstruction of distance distribution with SF-SVD.

**Figure 1.**
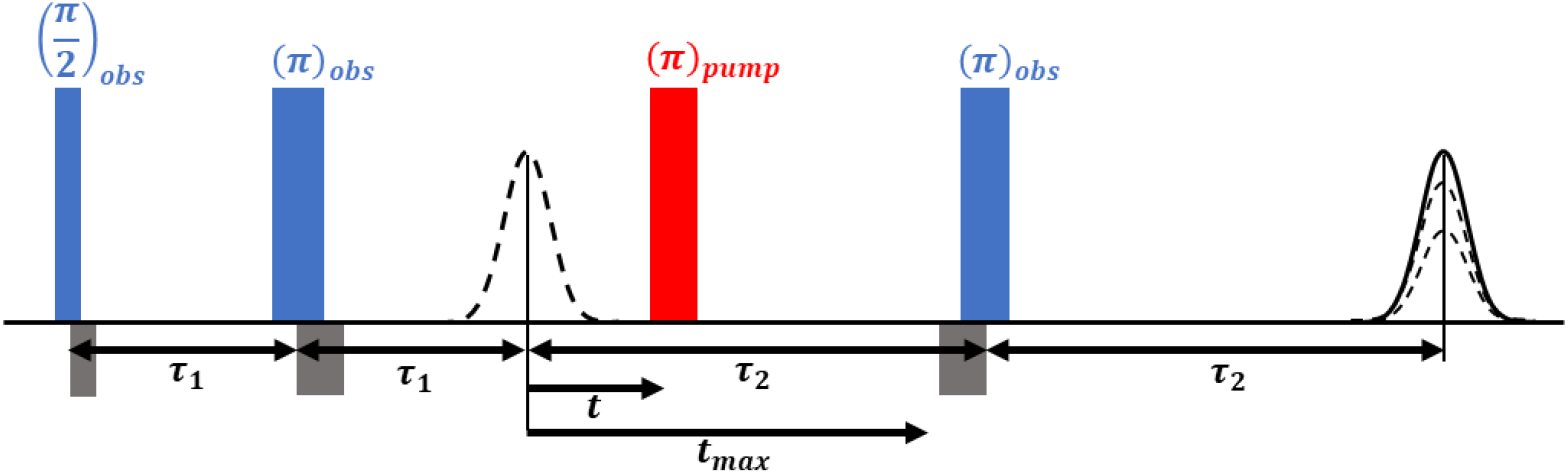
Four-pulse DEER sequence. Pump (red) and observer (blue) microwave pulses are used to selectively excite distinct spin populations, A and B. A two pulse Hahn echo is first formed by exciting A spins at the observer frequency. A pump pulse is subsequently applied to flip the B spins followed by a varying time delay, *t*, resulting in a modulation of the echo amplitude of A spins. At the delay, *τ*_2_, the echo is refocused by an additional pulse at the observer frequency. The DEER experimental trace, *V*(*t*), is the integral of the refocused echo as a function of pump pulse position, *t*.

**Figure 2.**
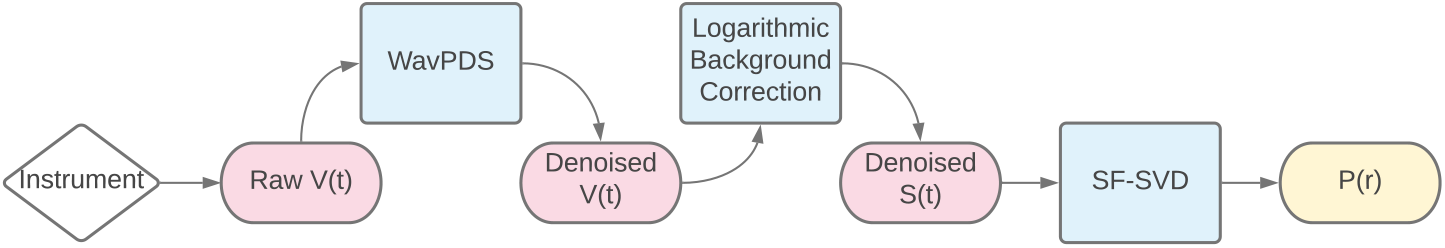
Data Processing workflow.

### 2.3 Theory of DEER Data Analysis

The objective of DEER is to quantify the distribution of dipolar interactions between pairs of spin labels that give rise to dipolar modulation in the time-domain signal. Due to the presence of an ensemble of sample conformations, the exact shape of the dipolar modulation depends on the distribution of distances, *P*(*r*), and yields a characteristic dipolar evolution function, *S*(*t*). The experimental time-domain signal derived from DEER contains the *S*(*t*) along with the background function in the following form^25,33^:

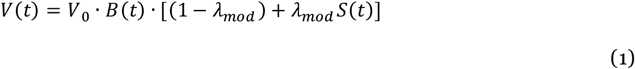

*V*_0_ is the overall amplitude of the unmodulated integrated echo (i.e. the maximum value of the signal). The modulation depth, *λ*_*mod*_, is a correction factor that compensates for the incomplete excitation of B spins by the pump pulse. The background function, *B*(*t*), results from intermolecular spin-spin interactions and is usually modeled as a stretched exponential decay^33–35^:

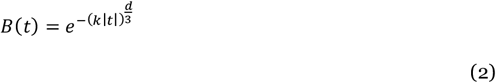

where *k* represents the background decay rate constant, which depends on the spin concentration and the pump efficiency, while *d* represents the dimensionality of the spatial distribution of the electron spins of the probes^8,36^. If the spin probe is homogeneously distributed in three dimensions, then *d* = 3. For samples of unknown dimensionality, *d* can be determined by fitting the above function to the experimental signal of singly labelled samples with the same properties as the doubly labelled sample^37,38^. *k* is determined by fitting the part of *V*(*t*) after dipolar modulation has dampened by equation 2 with selected *d*.

### 2.4 Background Correction

The background correction is performed using a subtraction method in the logarithmic domain. Fig. 3 shows the steps for background correction.

**Figure 3.**
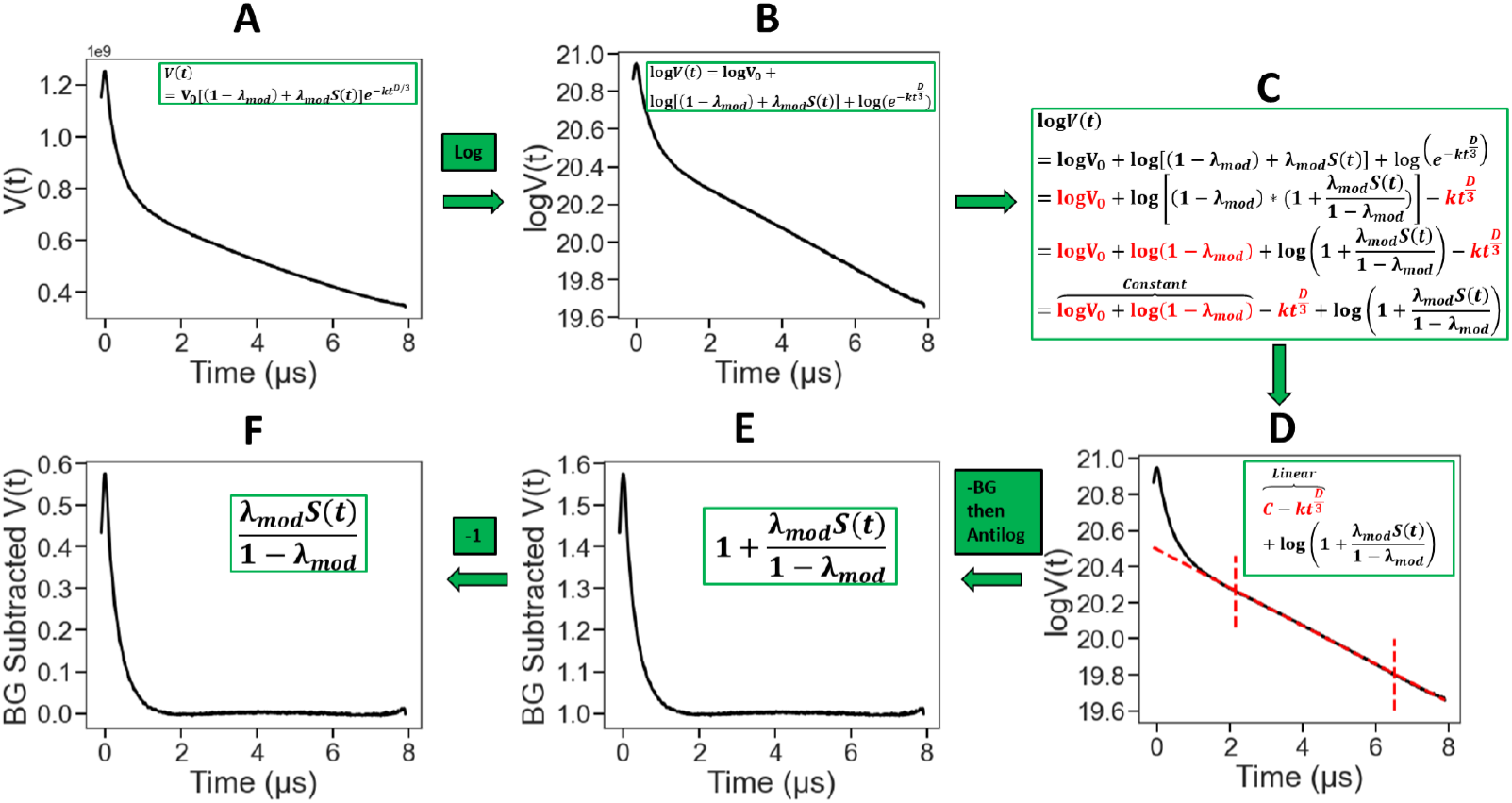
Background correction process using logarithmic approach to obtain *S*(*t*) from the measured *V*(*t*), after correcting for the background function, *B*(*t*) and the modulation depth, *λ*_*mod*_.

#### Steps A and B

From equations 1 and 2

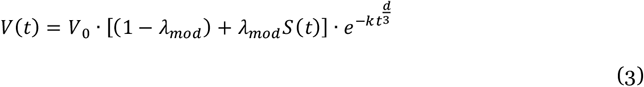

After taking the natural *log*: log(*A* · *B* · *C*) = log *A* + log *B* + log*C*

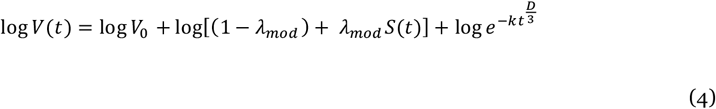

#### Step C

By rewriting log [(1 – *λ*_*mod*_) + *λ*_*mod*_ *S*(*t*) in the format: 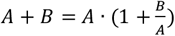, eq. 4 becomes:

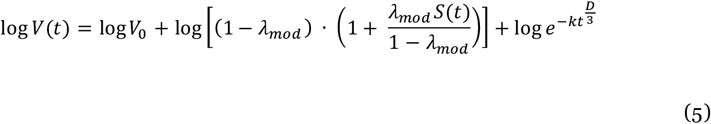

By rewriting log 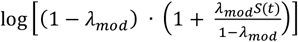 in the format: log(*A* · *B*) = log*A* + log *B*, and letting : log *e* ^*A*^ = *A*, we have from equation 5:

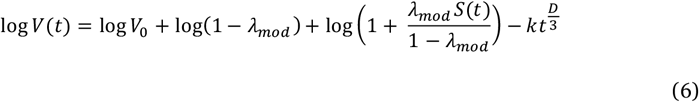

Rearranging equation 6:

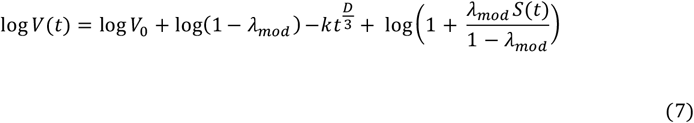

Let *Constant* = log *V*_0_ + log(1 − *λ*_*mod*_):

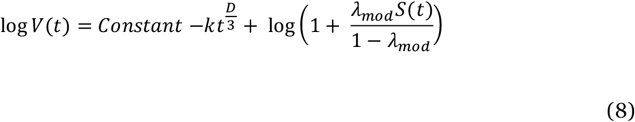

#### Steps D, E and F

After subtracting the linear part (cf. Fig. 4), (which includes background: 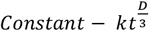) one has 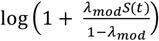 for which the antilog is 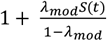. Subtract unity to yield:

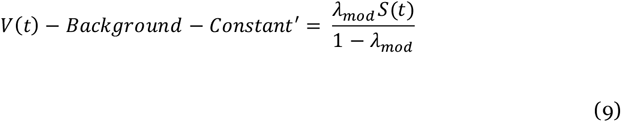

**Figure 4.**
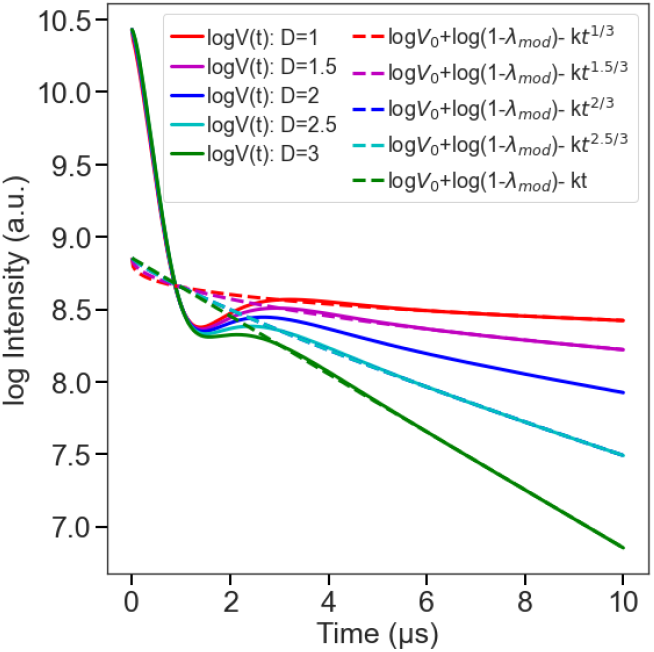
Background correction process using logarithmic approach revealing the linear part associated with the background function at different dimensionality (D) values.

Yielding *S*(*t*) after determining the modulation depth, *λ*_*mod*_.

It should be noted that for *λ*_*mod*_ = 1, equation 3 would be:

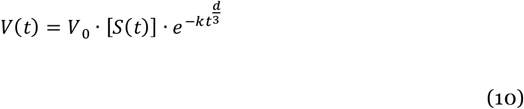

And hence, equation 9 would result as:

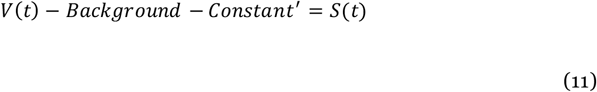

### 2.5 Inversion Problem

Continuing with *S*(*t*), this time-domain function can be expressed using the Fredholm integral of the first kind, where the problem is to find the function *P*(*r*) given the continuous kernel function *κ*(*t, r*) and the function *S*(*t*)^25^.

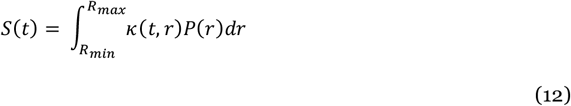

The kernel *κ*(*t, r*) represents the ensemble average over all dipolar couplings in the weak coupling limit in an isotropic mixture that results in a powder spectrum of orientation-dependent dipolar couplings for a given *r*. Since only the secular dipolar term is considered in the weak coupling limit, the dipolar kernel can be expressed as follows^25,34^:

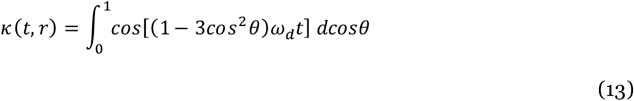

where 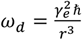 is the dipolar coupling constant, *γ*_*e*_ is the gyromagnetic ratio of each electron spin, and *ħ* is the reduced Planck constant.

The elements of eq 12 are discretized in the following way^25^:

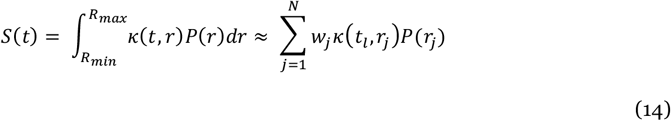

where 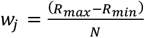 is the number of distance increments to be analyzed by SF-SVD, 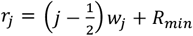 for *j* = 1, 2, …, *N* is the incremented distance variable, and 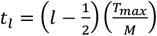 for *l* = 1, 2, …, *M* is the incremented time variable. The elements of eq 12 become *K*_*lj*_= *w*_*j*_ *κ*(*t*_*l*_, *r*_*j*_), *S*_*l*_ = *S*(*t*_*l*_), and *P*_*j*_ = *P*(*r*_*j*_), and hence ∑_*j*_ *P*_*j*_ *w*_*j*_ = 1. ^33^ Therefore, eq 12 can be presented as a system of linear algebraic equations in matrix notation:

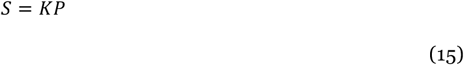

Because the kernel matrix *K* has a large condition number (meaning the ratio of the largest to the smallest singular value is substantially larger than 1), solving eq 15 to obtain *P*(*r*) is an ill-posed problem, meaning that computing the inverse of *K* will result in numerical errors and lead to unstable solutions for *P*(*r*) ^25,39^.

### 2.6 Srivastava-Freed Singular Value Decomposition (SF-SVD) Method^28,29^

#### 2.6.1 Singular Value Decomposition (SVD) of the Kernel Matrix

Instead of computing the inverse of the kernel matrix, the singular value decomposition of the Kernel matrix is required to obtain its pseudo-inverse, which can be used to extract stable *P*(*r*) from *S*(*t*). The kernel matrix *K* is an *M* × *N* matrix (*N* ≥ *M*), where the length of *M* is determined by the number of time-domain data increments in *S*(*t*) and the length of *N* is determined by the number of distance value increments chosen to resolve *P*(*r*). The *N* number of points can be equal or larger than the *M* number of points in *S*(*t*). *K* can be decomposed into the product of two square orthogonal matrices, *U* and *V*, and a diagonal matrix, *Σ*, written as *K* = *UΣV*^*T*25,28,29^:

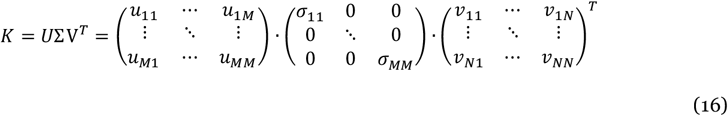

*U* has the dimension of *M* × *M, V* has the dimension of *N* × *N*, and *Σ* has the dimension of *M* × *N*. The matrix *V* contains the eigenvectors of *K*^*T*^*K* in its columns (also called right singular vectors), *U* contains the eigenvectors of *KK*^*T*^ in its columns (also called left singular vectors), and *Σ* contains the singular values, which are square roots of the non-zero eigenvalues of *K*^*T*^*K* or *KK*^*T*^ and acts as a scaling factor that allocates weight for each index element of *K. K* has the rank *k* (*k* < *M*), which is the number of non-zero singular values of *K* and is equivalent to the number of non-zero diagonal elements in *Σ*.^40,41^

#### 2.6.1 Point-wise Pseudo Inversion

The SVD of the kernel matrix allows it to be pseudo-inverted; therefore, *P*(*r*) can be obtained using the following relationship:

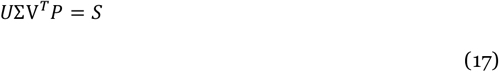

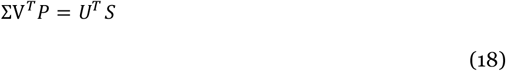

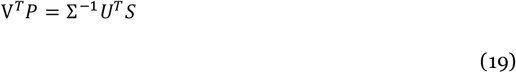

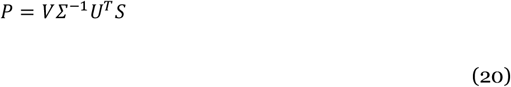

As *V* and *U* are orthogonal matrices, *V*^−1^ = *V*^*T*^ and *U*^−1^ = *U*^*T*^. *Σ*^−1^ is defined as,

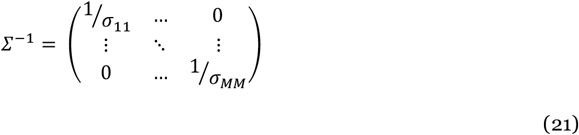

Therefore, *P* can be written as

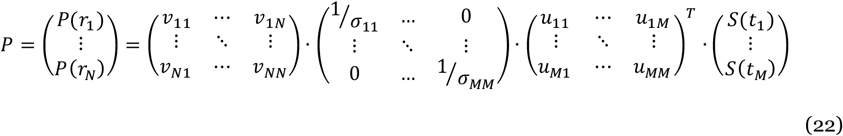

As one can see from eq 22, not all values of *V* contribute to each value of *P*, and hence each value *P*_*j*_ can be written in dependence of *V* as follows:

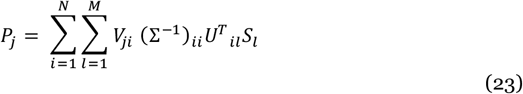

where *N* is the dimension of the discrete values of *P* with the index, *j*, representing distance *r*, and *M* of *S* with *l*, representing time *t*, and *i* is the index representing the singular values, *σ*_*i*_ ‘*s*, along the diagonal of *Σ*^−1^.

To find the optimal solution for *P*_*j*_ by SF-SVD, an individual singular value cutoff, *k*_*j*_, can be chosen for each distance or distance range. This is one of the key features of SF-SVD. The singular value contribution to *P*_*j*_ can now be written as (*SVC*_*j*_)_*i*_, a vector summed over index *i* from 1 to the cut off, *k*_*j*_, as shown below:

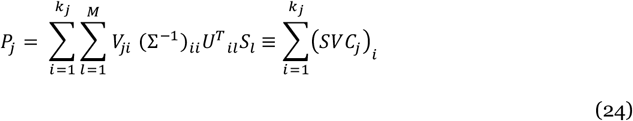

The singular value cutoff that varies for each distance increment can now be represented as *k*_*j*_, and can be determined by a modified Picard condition as follows:

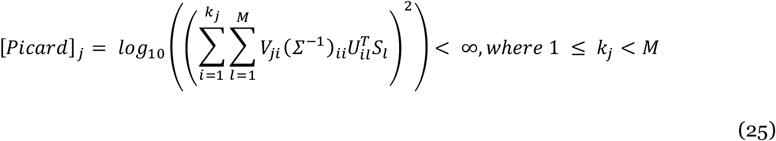

As explained in the previous section, *k*_*j*_ is the definite divergence point that occurs when *k*_*j* +1_ singular value contribution (*SVC*_*j*_)_*i*_ is added to the sum.

In a special case, where all the SVC cutoffs are same (i.e. *k*_1_ = *k*_2_ =.. *k*_*j*_. = *k*_*N*_), the exact solution can be obtained by a single singular value cut-off *k* and can be represented in the following way:

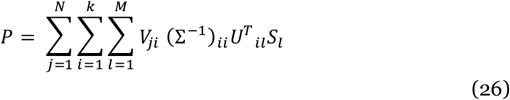

The *k* cut-off value can be obtained using the Picard plot as follows:

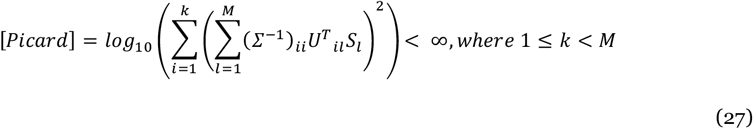

In the above mentioned case, the cut-off *k* value also reflects the rank of matrix and according to a theorem^41^, there exists an exact solution for *P*, if *S* is orthogonal to the *M* − *k* left singular vectors of 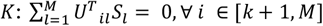. In other words, all *Σ* with *k* < *i* ≤ *N* are zeros, and therefore, it must be neglected. Fig. 5C shows *k* is determined as the point before Picard plot diverges from the flat region on the plot. Note that in Fig. 5C the value has converged well before it diverges.

**Figure 5.**
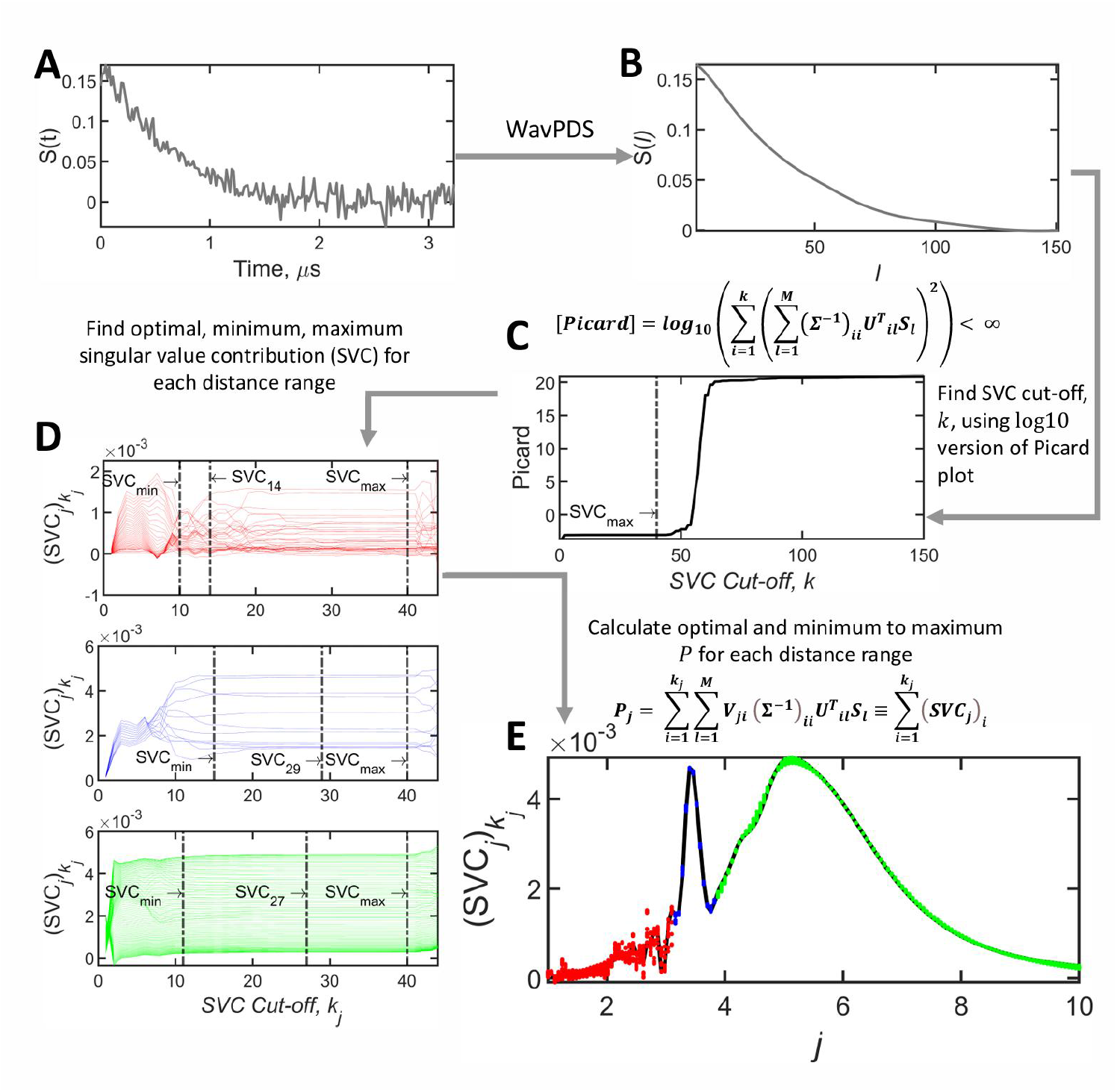
(A) Noisy background-corrected DEER signal from tau 187 WT fibril labeled at site 291-322 (c.f. Fig. 6c). (B) Denoised background-corrected signal using WavPDS. (C) The Picard plot 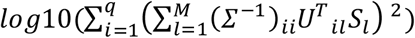 versus the SVC cut-off *k*, showing after *i* = 40, (the Picard value is denoted as *SVC*_*max*_ at the maximum cut off index, *k*, for singular value contributions), the solution becomes unstable. (D) The SVC contributions for the index j, (*SVC*_*j*_)_*i*_, for 3 different distance regions (i.e. spanning 3 different j ranges), indicated in red (1-3.1 nm), blue (3.16 – 3.82 nm), and green (3.88 – 10 nm), are plotted against the SVC cut-off *k*_*j*_. Instead of showing the plot for the entire range of *k*_*j*_ as in (C), in (D) the (*SVC*_*j*_)_*i*_, is plotted only in the range between the minimum and maximum cut off index *k*_*j*_, including the optimal cut off index, for a series of *j* indices, (E) The 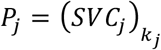 values are plotted vs the index *j*, with the 3 distance regions reconstructed using different optimum SVC values are plotted in red, blue, and green, making up the optimal distance distribution plotted in black.

#### 2.6.2 SF-SVD: Convergence and Uncertainty Determination

The SF-SVD method obviates the need for a regularization parameter [28]. Instead, it obtains each *P*(*r*_*j*_) amplitude for a given *j* as a function of the (*SVC*)_*i*_ value. The existence of an exact solution that is intrinsic to the theorem^41^ implies that each *P*(*r*_*j*_) must converge at a certain value for *i* = *k*_*j*_ as (*SVC*_*j*_)_*i*_ contributions are added from *i* = 1 to *k*_*j*_. This convergent value does not change until the Picard condition is met leading to divergence. When some random noise is added in the convergence region, there is some fluctuation in *P*(*r*_*j*_) and the Picard condition occurs a little sooner vs. (*SVC*_*j*_)_*i*_ but the convergence remains [28]. This is illustrated in Fig. 5D for different distance ranges segmented in three *j* ranges. Of course, too much noise can obscure these important features, so in such cases it is necessary to first denoise the data by WavPDS. This has been shown to be true for many cases^4,6,30,31^. The fluctuation obtained for *P*(*r*_*j*_) over the convergent range of the (*SVC*_*j*_)_*i*_ (from SVC_min_ to SVC_max_) due to the residual noise is taken as the uncertainty in that value, while the mean yields the value of *P*(*r*_*j*_). As we have already pointed out, the convergence region and the fluctuations differ for different values of *r*, and hence the index *j*, as illustrated in Fig. 5D. The distance distribution obtained from such convergence is shown in Fig. 5E.

## 3. RESULTS

Here, we examine the ability of WavPDS and SF-SVD to reconstruct the desired probability distribution of distances, *P*(*r*), under a variety of conditions, such as different noise levels, dipolar evolution times, and experimental PDS signal, *V*(*t*), from different sample types. We chose different sets of molecular systems that give rise to distinct and narrow, uniformly broad, or narrow-broad mixed *P*(*r*) shapes. Specifically, we chose stiff end-to-end spin labeled molecules (Fig. 6a), oligo(para-phenylene ethynylene), with unimodal narrow distributions centered around 2.8 nm, 4.1 nm and 5.4 nm, to represent systems with distinct and narrow distributions^42,43^. In this paper, we named it Ruler 1, 3 and 5 for n=1, 3, 5 chains. We chose a set of end-to-end spin labeled polyethylene glycol (PEG) polymers, comprising of repeating units of [-CH_2_-O-CH_2_-] terminated by [-OH], with four different molecular weights between 220 and 1600 Da (Fig. 6b). PEG is disordered and semi-flexible in water, which gives rise to an ensemble of conformations leading to broad *P*(*r*) ; therefore, it is a good model system for uniformly broad distance distributions. Next, we apply this method to the target samples, the intrinsically disordered protein (IDP) tau and its fibrillar state (Fig. 6c).

**Figure 6.**
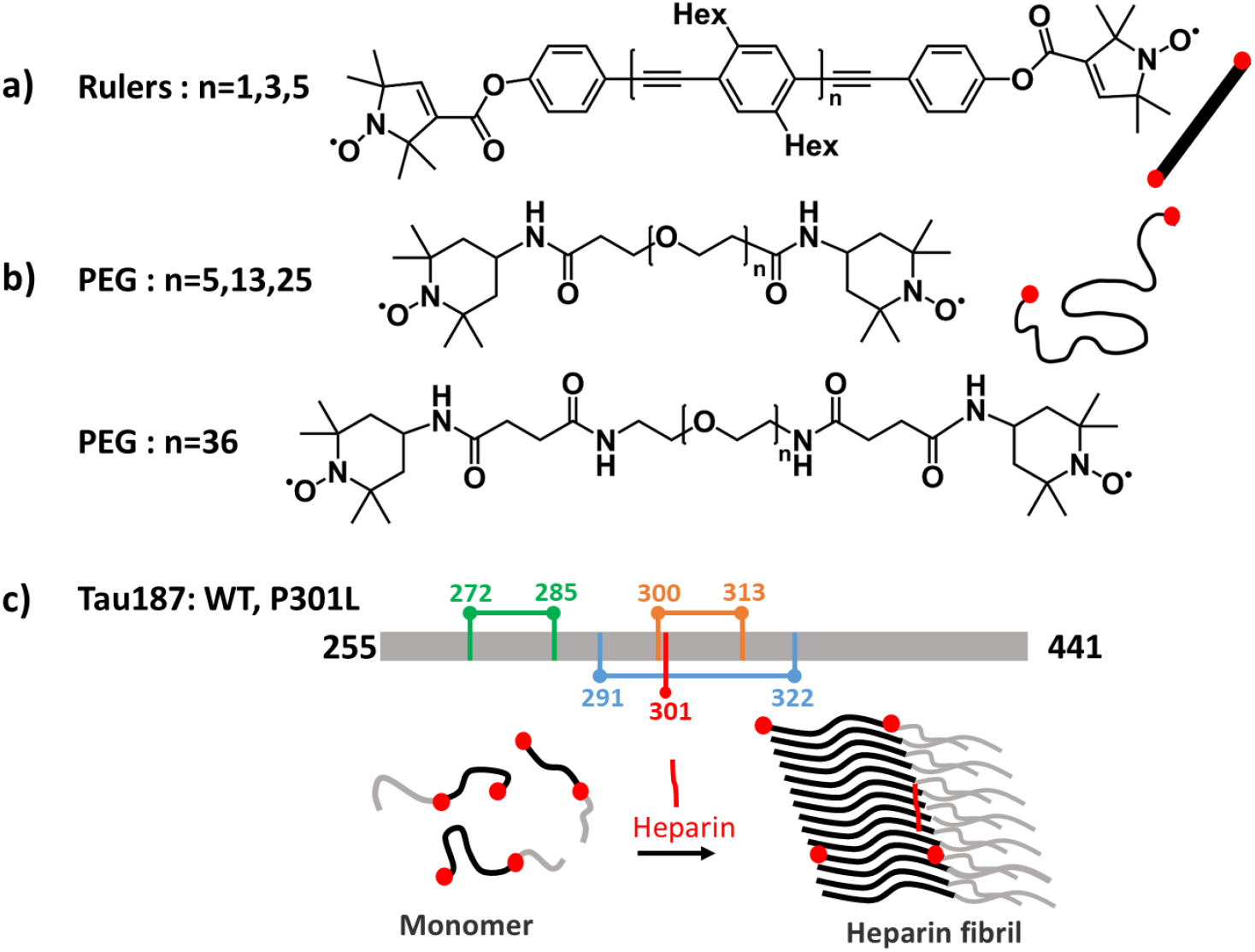
Molecular systems analyzed by DEER and SF-SVD: a) DEER ruler (oligo(para-phenylene ethynylene)) chains with 1,3, and 5 monomer inserted, b) PEG (polyethylene glycol) chains with 5, 13, 25, and 36 monomer inserted, and c) tau187 P301L (with mutation of Proline to Leucine at residue 301 in red) monomer and fibril that is spin labeled at either residues 272-285 (green) or 300-313 (orange), and tau 187 WT monomer and fibril that is spin labeled at 291-322 (blue).

N0te that for comparisons of the SF-SVD results with the TIKR methods, we use the DEERLab^26^ and LongDistance^27^ software. The results are not compared with PD_TKN and MEM software^25,44^, as it was retired after the introduction of SF-SVD method by Srivastava and Freed. The results between them are compared in our previous work^28^. GLAADvu^20,21^ and DEERNet^22,23^ are not used for the reasons explained in the introduction.

Note that for simplicity of notation, we denote the background corrected experimental DEER signal as dipolar signal, *S*(*t*), for the remainder of the paper.

### 3.1 Reconstruct distance distribution from low SNR data by denoising

First, we set out to test the ability of WavPDS and SF-SVD to reconstruct the desired *P*(*r*) distribution from low SNR dipolar signals, *S*(*t*). For this, we used the stiff Ruler 1, 3 and 5 molecules with known distances between nitroxide spin labels positioned at both ends (cf. Fig. 6a). The data from three rulers with well-defined distributions centered at 2.8nm, 4.1 nm and 5.4nm, respectively, are collected with different scan numbers for signal averaging to generate low and high SNR data (Fig. 7, left to right, respectively). An evolution time of 8 *µs* was selected to maximize the accuracy for the reconstruction of the distance distribution. Denoising was applied to each of them to obtain high SNR, after which SF-SVD was used to reconstruct *P*(*r*). As can be seen from Fig. 7, the mean value of the unimodal *P*(*r*) distributions obtained from n=1, 10 and 428 scans for the 2.1nm ruler is identical and at the expected distance; a similar observation is made for the *P*(*r*) of the 4.1 nm and 5.4 nm ruler. Apart from the mean value, the shape of *P*(*r*) is also reliably recovered. These results showcase that SF-SVD coupled with WavPDS can be readily applied to reconstruct well-defined distance distributions from experimental PDS data at low SNR data acquired with a low number of scans.

**Figure 7.**
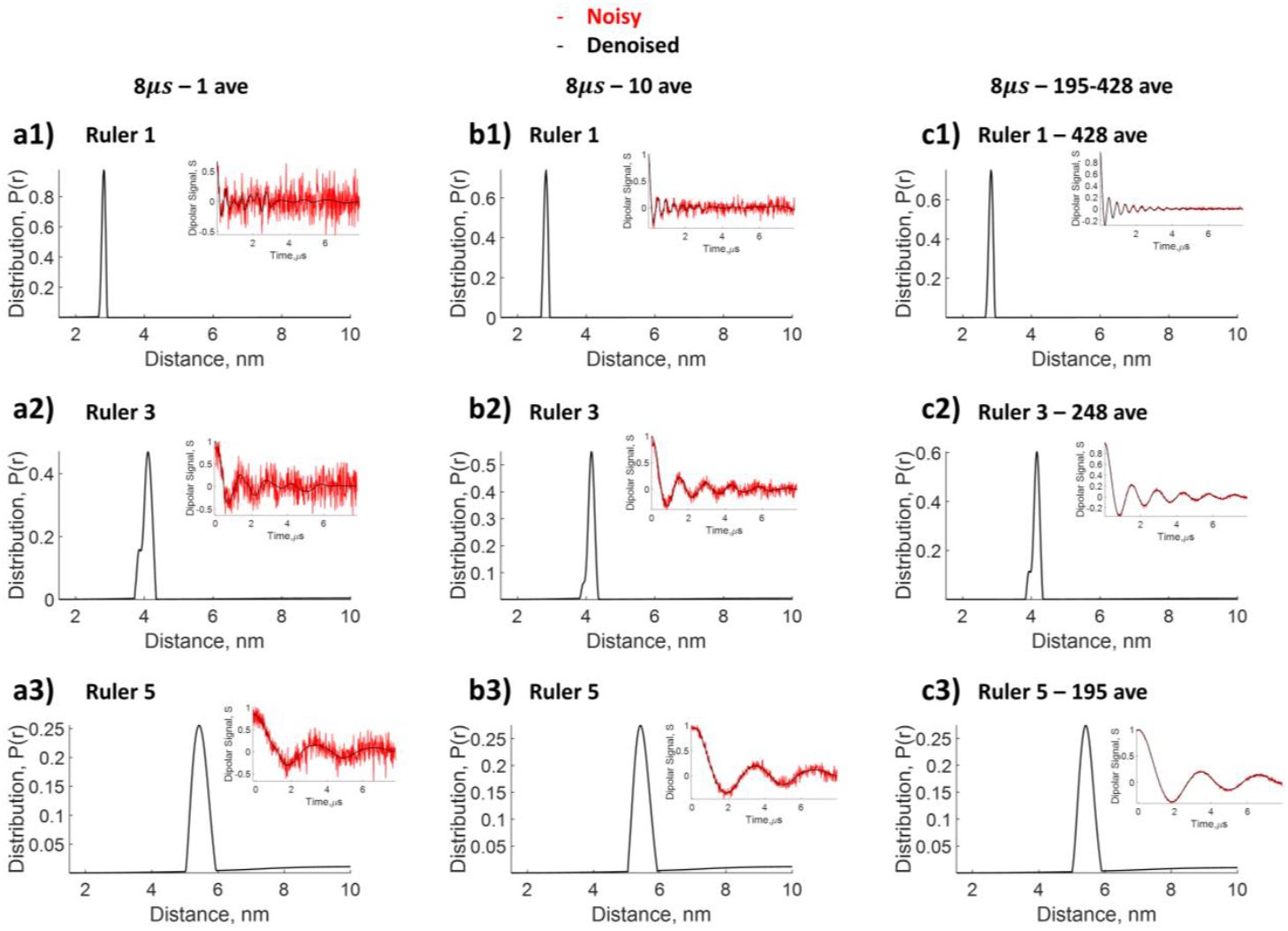
Comparison of distance distributions for DEER rulers reconstructed from dipolar signal different number of scans, but with the same 8 *µs* evolution time. n=1 ruler with expected distance of 2.8 *nm* **a1)** 1 average, **b1)** 10 average, **c1)** 428 average; n=3 ruler with expected distance of at 4.1*nm* **a2)** 1 average, **b2)** 10 average, **c2)** 248 average; and n=5 ruler with expected distance at 5.4*nm* **a3)** 1 average, **b3)** 10 average, **c3)** 195 average. SF-SVD after denoising is used to reconstruct distances.

### 3.2 Reconstruct distance distributions from dipolar time domain data with different evolution times

Longer evolution times to acquire the time-domain dipolar signal enhance the resolution of the reconstructed distance distributions. However, acquiring DEER data with long evolution times is time-consuming, and hence, often a compromise is found between the number of scans and the evolution time for DEER. The choice of longer evolution time often comes with the challenge of greater noise, i.e. low SNR, obviating any gain associated with the longer evolution times. Data from three rulers (n=1, n=3, and n=5) were acquired at three different evolution times (3*µs*, 5*µs* and 8*µs*) with the same number of scans of 10 (i.e. same data collection time). This choice yielded high SNR at short evolution time and low SNR at long evolution time. The WavPDS and SF-SVD methods (cf. Section 2) were subsequently applied to each dataset. As shown in Fig. 8, the results from the ruler data collected at different evolution times yield the same mean distance, but varied shapes for *P*(*r*). The results for the n=3 and n=5 ruler show that the shape narrows, i.e. is better resolved with an increase in the dipolar evolution time from 3 *µs* to 5 *µs*, and further to 8 *µs*. For the shortest n=1 (i.e 2.8 nm) ruler, *P*(*r*) narrows when lengthening the dipolar evolution time from 3*µs* to 5*µs*, but remains unchanged with the longer evolution time of 8*µs*. Hence, the 5 *µs* evolution time is sufficient to accurately reconstruct the shape for *P*(*r*) for this short distance, but not for the longer 4.1 nm and 5.4 nm distances. This exercise demonstrates the capability of WavPDS and SF-SVD to reconstruct well-resolved distance distributions at longer evolution times, even from noisy data. Hence, denoising by WavPDS offers the opportunity to acquire dipolar time-domain data at longer evolution times, yielding better resolution, even with limited SNR such as those of DEER data with only one scan average (cf. Fig. 7).

**Figure 8.**
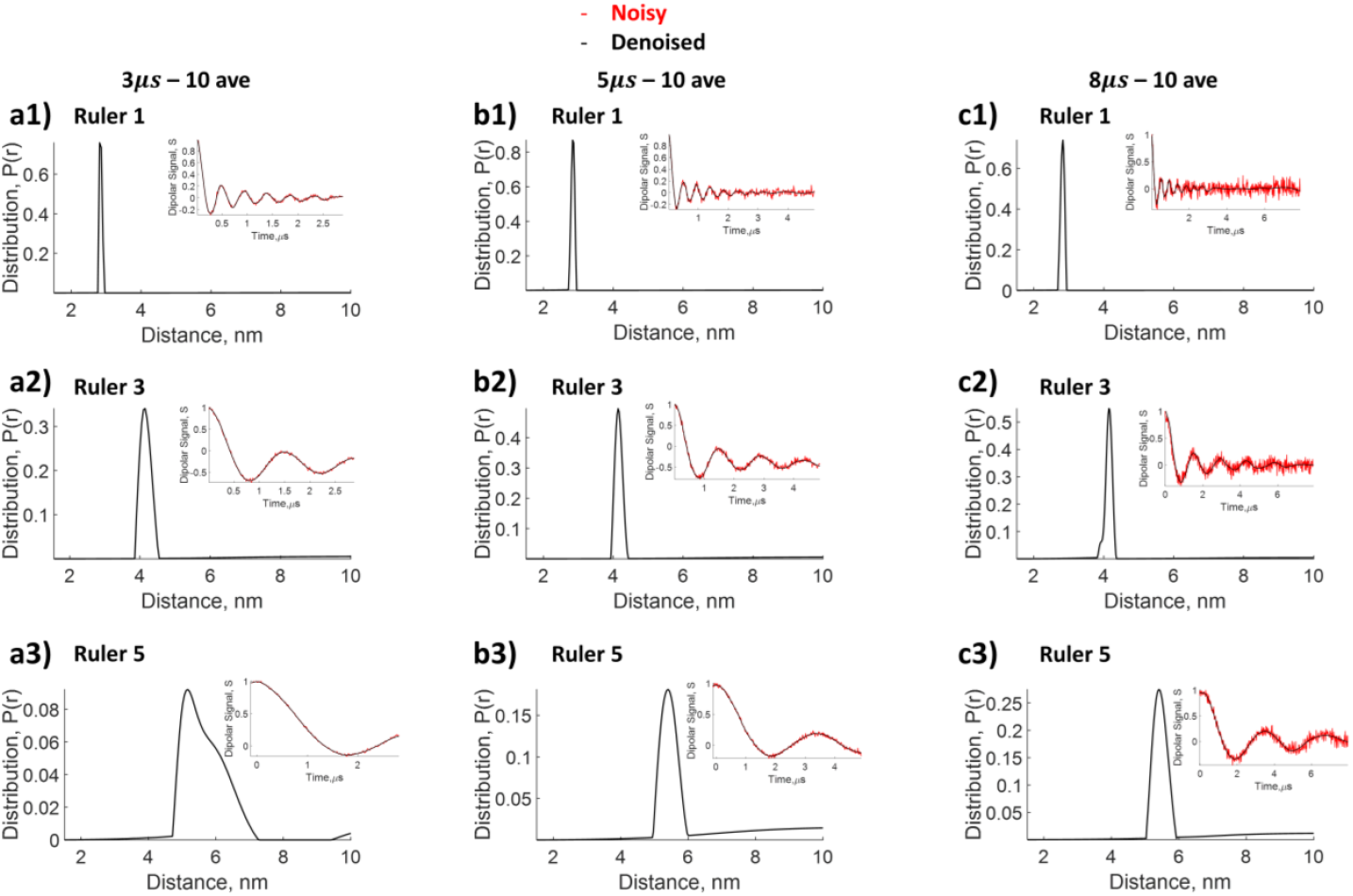
Comparison of distance distribution reconstructed from dipolar signal with different evolution times, but the same number of scans of 10. N=1 ruler with expected distance of 2.8 *nm* **a1)** 2*µs*, **b1)** 5*µs*, **c1)** 8*µs*; n=3 ruler with expected distance of 4.1*nm* **a2)** 2*µs*, **b2)** 5*µs*, **c2)** 8*µs*; and n=5 ruler with expected distance of 5.4*nm* **a3)** 2*µs*, **b3)** 5*µs*, **c3)** 8*µs*. SF-SVD after denoising is used to reconstruct distances.

### 3.3 Comparison of SF-SVD and Tikhonov Regularization for the *P*(*r*) Shape

It is a challenge to reliably determine the shape for *P*(*r*) of intra-molecular distances of an IDP. We first show the effectiveness of SF-SVD in generating mean and shape information for a well-defined *P*(*r*) of the n=1 and n=3 rulers (cf. Fig. 9 a4 and b4), without the need to select any optimal parameters. The results are compared with TIKR with choices of *λ* influencing the distribution outcome. SF-SVD is applied after WavPDS denoising, whereas TIKR is applied to both original noisy and denoised data for comparison. As can be seen from Fig. 9, the SF-SVD reliably recovers both the mean and the narrow distance distribution shape (rulers are rigid molecules and have narrow *P*(*r*)^′^*s* shapes). TIKR can reliably yield mean distances, both for noisy and denoised cases. However, as expected, the shape of *P*(*r*) derived from TIKR depends on the choice of regularization parameter *λ*, making it more ambiguous to use TIKR for shape determination. The values for the regularization parameter *λ* were varied from 20 to 0.1 for TIKR, shown in Figure 9 (a2, b2, a3, b3). As can be seen, the choice of *λ* yields a large variation in distance distributions, in particular the width, that gives rise to uncertainty in the shape determination. There are several ways to determine optimal *λ* values using the regularization parameter selection criterion such as the L-curve^24,25,27^ and the Akaike Information Criterion^33^, among others (cf. Table S1). These methods use distinct objective criteria to find optimal values and can generate different optimal *λ* values. This can be seen from Fig. S1 and Table S1 (cf. Supporting Information) for the n=1 and n=3 rulers, where the optimal *λ* values generated from the various criterion yield different values, ranging from 0.0001 to 15082 for the n=1 ruler and 0.0003 to 67538 for the n=3 ruler. It is, however, worth mentioning that many optimization parameters yield values ranging from 0.02 to 0.04 for ruler n=1 and 0.4 to 0.8 for n=3, suggesting that the distance distributions under those *λ* values are more reliable. On the other hand, the mean distance value remains unchanged with the choice of *λ*. The TIKR method can yield the desired shape in a narrow distance distribution, provided that the appropriate regularization parameter is selected either by human judgement and/or *a priori* information.

**Figure 9.**
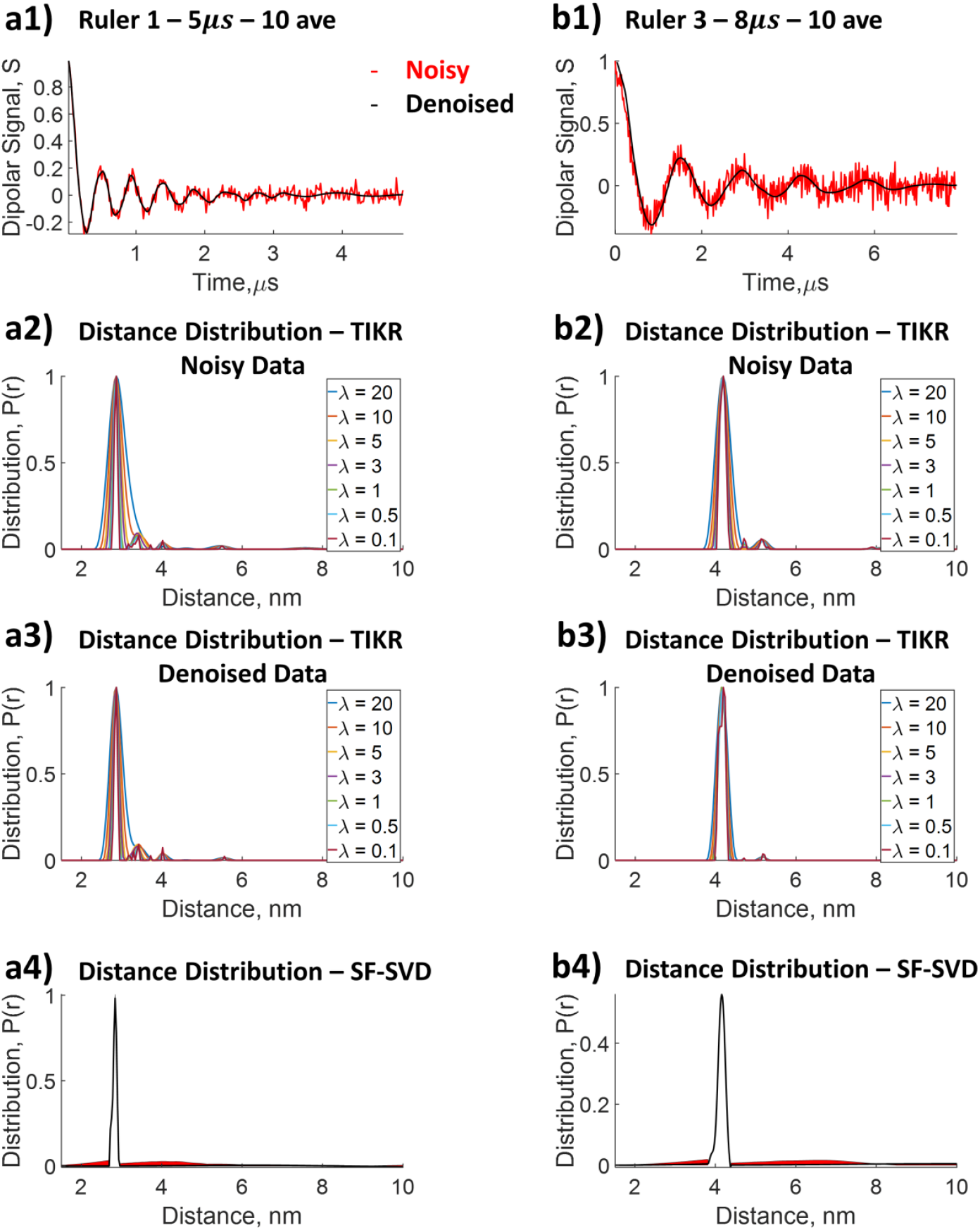
Comparison of distance distribution reconstructed from dipolar signal by Tikhonov regularization with different regularization parameter values, *λ*, and that by SF-SVD using denoised dipolar signal. **a1)** n=1 ruler with dipolar signal collected at 5*µs* evolution time at 10 averages; **a2)** *P*(*r*) from Tikhonov regularization for n=1 ruler; **a3)** *P*(*r*) from Tikhonov regularization of denoised data for 2.8*nm* ruler; **a4)** *P*(*r*) from SF-SVD for n=1 ruler; **b1)** n=3 ruler with dipolar signal collected at 5*µs* evolution time at 10 averages; **b2)** *P*(*r*) from Tikhonov regularization for n=3 ruler; **b3)** *P*(*r*) from Tikhonov regularization of denoised data for 2.8*nm* ruler; **b4)** *P*(*r*) from SF-SVD for n=3 ruler.

SF-SVD, on the other hand, yields an appropriately narrow distance distribution for n=1 and n=3 rulers, without the need to rely on the choice of *λ*. To recapitulate: the solution is obtained by finding the convergent solution independently for each distance or distance range w.r.t. singular value contributions to the solution. The convergent values are obtained from the procedure illustrated in Fig. 5. Hence, the shape determination is accurate and reliable, and one can probe the influence of individual singular values at a given distance.

Here, WavPDS aids SF-SVD by denoising of the time-domain data to reduce the uncertainty in the solution. TIKR also benefits from WavPDS, and the P(r) shape is better resolved after denoising. Even so, TIKR only reliably yields information about the mean distance.

### 3.4 Reconstruction of Broad Distributions of End-to-End Labeled PEG

We next show the result of the synthetic polymer PEG (polyethylene glycol) chains with 5, 13, 25, and 36 monomers (220 to 2600 Da) derived from SF-SVD (cf. Fig. 10 row 1), LongDistance (cf. Fig. 10 row 2), DD Gaussian (cf. Fig. 10 row 3), DeerLab (cf. Fig. 10 row 4), and DEERNet (cf. Fig. 10 row 5). In Fig. 10 row 1, a single broad shape is obtained using SF-SVD, where the SVC contributions and the singular value cut-off ranges around 0.16, reflecting a well-resolved distribution. This is important in demonstrating SF-SVD’s ability to differentiate a broad *P*(*r*), due to disordered structures, from *P*(*r*) that contain multiple distinct peaks due to partially ordered structures in heterogeneous samples. One can also see in Fig. 10 row 1 the asymmetry in the *P*(*r*) shape, associated with the PEG data. The *P*(*r*) derived from TIKR^45^ was recently published and analyzed using model-free fitting by LongDistances^27^ with a *λ* of 30. The broader shape and smoothed distributions in TIKR are due to choosing a regularization parameter, *λ*, that is too high (i.e. 30), and is often the case for all the broad distributions. SF-SVD does not suffer from such consequences because each distance point is independently optimized; therefore, a large regularization parameter is not needed to stabilize one region of the distribution, often leading to over smoothing of other regions. It is most evident in the 5 mer, where the distance distribution obtained by TIKR (c.f. Fig. 10 a2) is broader than by SF-SVD (c.f. Fig. 10 a1). In Fig. 10 row 3, the *P*(*r*) derived using gaussian model fitting using DD MATLAB software shows the expected broad distributions of PEG but with subpopulations. The subpopulation in longer PEG exists because the number of gaussians used for fitting was based on the least reduced chi square, which leads to fitting more than 1 gaussian. In Fig. 10 row 4, the *P*(*r*) derived using DeerLab python software shows the expected broad distribution of PEG with sharp spikes. DeerLab uses a one-step fitting where the background and *P*(*r*) parameters are simultaneously fitted with the regularization parameter automatically picked using the AIC criterion^26^. Therefore, the sharp spikes can be either from slight errors in background fitting or too small of a regularization parameter, which is partially dictated by the SNR of the time domain signal. The time domain signal of this PEG dataset has high SNR; therefore, the AIC criterion chose a small regularization parameter and less smooth result than that of LongDistance. With the same background correction as LongDistance and a higher regularization parameter, the DeerLab result will be comparable to that of LongDistance. Lastly, in Fig. 10 row 5, the *P*(*r*) derived using DEERNet MATLAB software shows the expected broad distribution of PEG with similar shape to the *P*(*r*) derived from SF-SVD and LongDistance. The P(r) for PEG 36 (c.f. Fig. 10d5) has a minor population around 6 nm that is not seen in the SF-SVD or the LongDistance result. Regardless of the spikes or subpopulations in the *P*(*r*), all five methods resulted in similar overall *P*(*r*) shape and mean distance.

**Figure 10.**
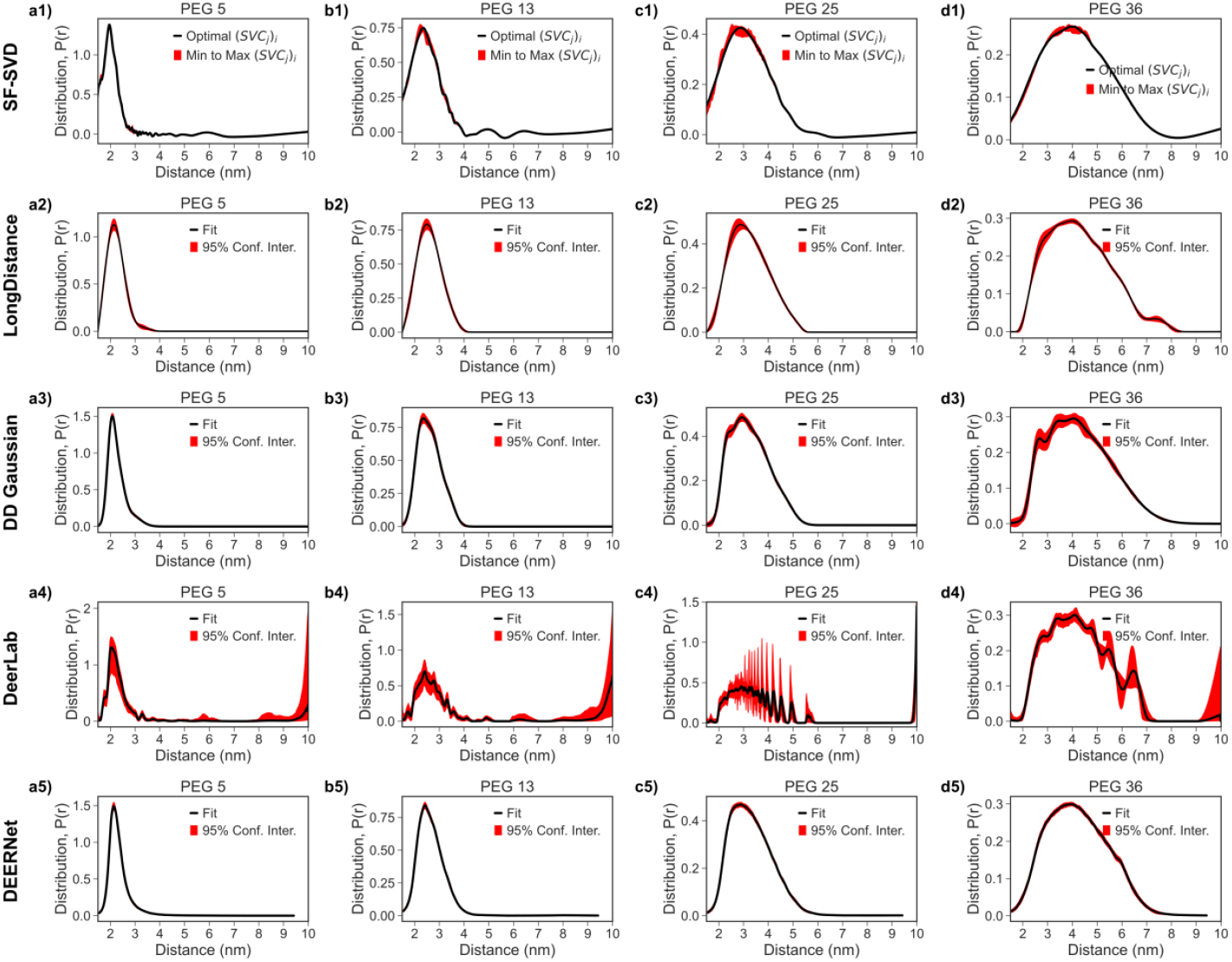
Comparison of Distance Distribution *P*(*r*) obtained from PEG DEER processed with the SF-SVD, LongDistance, DeerLab, DEERNet, and DD Gaussian methods. Column 1 **(a1-a5)** represents the end-to-end labeled PEG 5. Column 2 **(b1-b5)** represents the end-to-end labeled PEG 13. Column 3 **(c1-c5)** represents the end-to-end labeled PEG 25. Column 4 **(d1-d5)** represents the end-to-end labeled PEG 36. Row 1 **(a1, b1, c1, d1)** represents *P*(*r*) from the SF-SVD method. Row 2 **(a2, b2, c2, d2)** represents *P*(*r*) from the LongDistance method. Row 3 **(a3, b3, c3, d3)** represents *P*(*r*) from the DD Gaussian method. Row 4 **(a4, b4, c4, d4)** represents *P*(*r*) n from the DeerLab method. Row 5 **(a5, b5, c5, d5)** represents *P*(*r*) from the DEERNet method.

### 3.5 Reconstruction of More Complex Distributions: Examples from Tau

The power of the SF-SVD method is best illustrated in our studies of the intrinsically disordered protein (IDP) tau. IDPs can be thought of as complex biopolymers that are mainly deprived of secondary structures, but still can engage in specific intramolecular interactions^47^. Tau is present in the human brain where it regulates microtubule activity. Under pathological conditions, the microtubule binding tau protein forms β-structured amyloid aggregates that are involved in several neurodegenerative diseases including Alzheimer’s disease.^48^ Here we study a 187 amino acid-long, N-terminal truncated human 4R tau that contains residues 255-441, referred to as tau187. We examined both the wild type and mutant P301L tau187 with the residue 301 mutated from Proline to Leucine. We measured three different intramolecular distances by placing the labels either at residues 272-285 (13 amino acids apart), at residues 300 and 313 (also 13 amino acids apart), or at residues 291-322 (31 amino acids apart) (cf. Fig. 6c). For tau in its monomeric state all three intramolecular distances 272-285 (Fig. 11a1), 300-313 (Fig. 11a2), and 291-322 (Fig. 11a3) distances showed relatively featureless, broad, *P*(*r*), consistent with tau in its monomeric form being an IDP. The *P*(*r*) has max peak at 2.5 nm and spans 1.5-5.5 nm for the intramolecular 272-285 distances, max peak at 2.6 nm and spans 1.5-6 nm for the 300-313 distances and max peak at 3.6 nm and spans 1.5-8 nm for the 291-322 distances. Next, DEER measurements were performed on fibrillar tau187, induced with heparin, of the same sets of spin labeled tau187 (cf. Fig. 6c). Note that in all cases 10% of doubly spin labelled protein was diluted with 90% of unlabeled protein to ensure that intra, and not inter, molecular distances are measured. In contrast to monomeric tau187, we see that the distributions (cf. Fig. 11b) of the aggregated sample contains several components that could reflect different aggregate conformations. We observe at least three populations centered at 2.7 nm, 3.6 nm, and 4.2 nm for the 272-285 fibril (Fig. 11b1), three populations centered at 2.9 nm, 3.9 nm, and 5.5 nm for the 300-313 fibril (Fig. 11b2), and two populations centered at 3.6 nm and 5.3 nm for the 291-322 fibril (Fig. 11b3), (although the second population in the *P*(*r*) of 291-322 fibril appears rather broad and might encompass more than one conformation). Furthermore, the peak and the span of distances for each doubly spin labeled tau187 shifts towards larger values when tau187 transforms from a monomeric to a fibrillar state. In other words, tau187 is extended across sites 272-285, 300-313 and 292-322 when it incorporates into fibrils. This conformational extension for heparin-induced tau187 fibrils across sites 272-285 and 300-313 is consistent with the fibril structure (PDB 6qjh, 6qjm, and 6qjp) according to cryogenic electron microscopy (Cryo-EM) of the snake, twister, and jagged form of heparin-induced fibrils^46^. The observation of distance extension across the two natural cysteine sites, 292-322, cannot be easily rationalized by Cryo-EM structures given that none of the tau fibril structures encompass the 292 sites. However, it is shown in the literature^49,50^ that tau187 across 292-322 in the monomer form may populate a “folded-over” conformation and is intrinsically disordered, while site 292 is suggested to be in the fuzzy core of the tau fibrils (illustrated in Fig.6c and Fig. 11 as grey wiggles). Hence, the extension of the 292-322 distance is consistent with the organization of tau187 from an IDP to an organized fibril. The results demonstrate the viability of SF-SVD to find user-independent solutions to broad and featureless *P*(*r*) of the monomeric tau187, and *P*(*r*) of tau187 fibrils that contains both a narrow and a broad distance distribution population.

**Figure 11.**
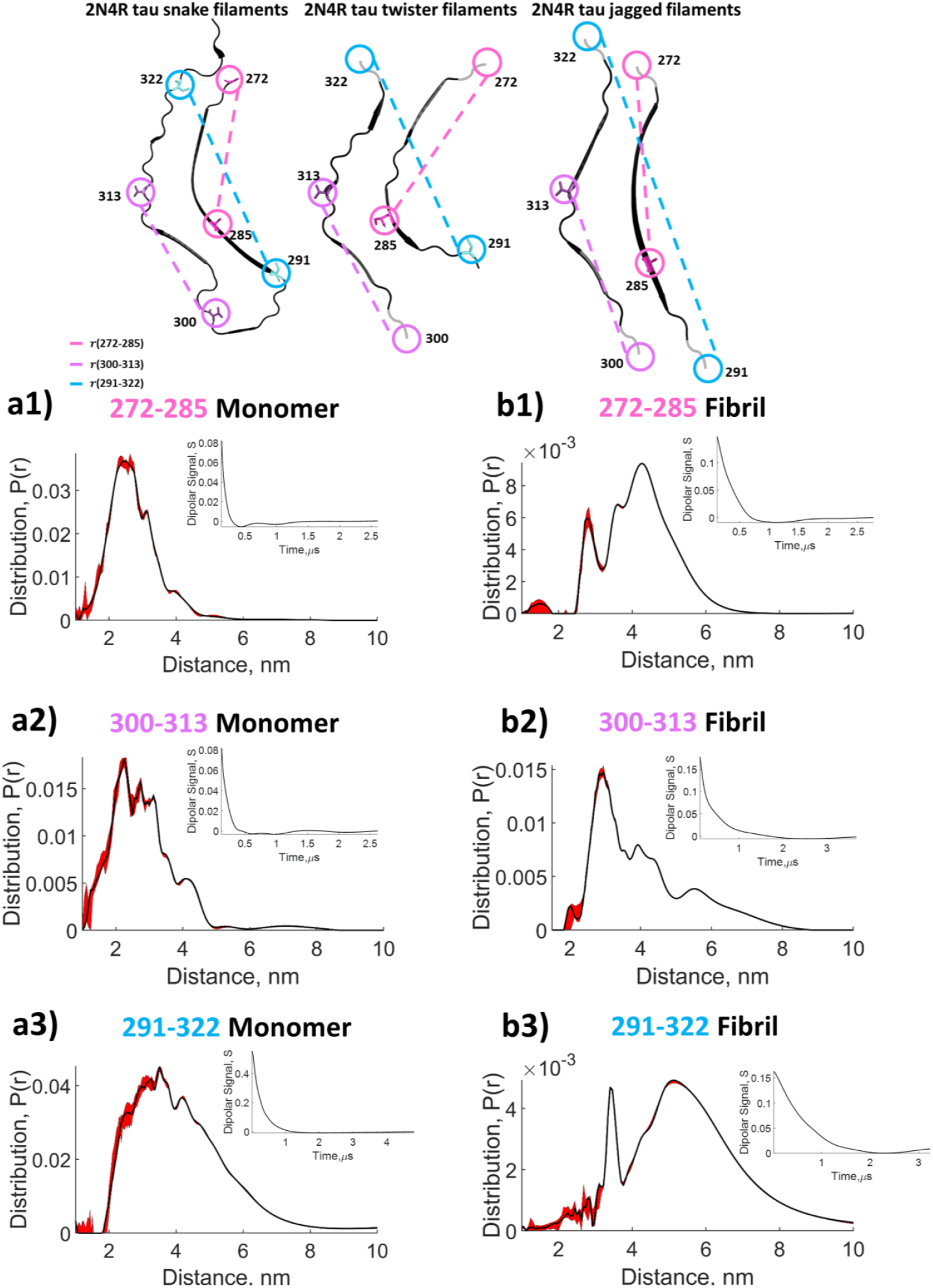
Top panel shows the schematic view (from PDB of structures solved by W. Zhang, et al^46^) of the 3 dominant structure of heparin induced 2N4R tau fibril. The grey circle and wiggle line are drawn to indicate residue not included in the cryo-EM analysis but are labeled in our tau187 sample. Distance distribution obtained with SF-SVD for the IDP tau187 monomer and amyloid fibril: *P*(*r*) of tau187-P301L labelled at residues 272-285 (pink) measured in its **a1)** monomeric state and **b1)** aggregated form; *P*(*r*) of tau187-P301L labelled at residues 300-313 (lilac) measured in its **a2)** monomeric state and **b2)** aggregated form; *P*(*r*) of tau187-WT fibril labelled at residues 291-322 (blue) measured in its **a3)** monomeric state and **b3)** aggregated form. The red area around the curve represents the uncertainty. The denoised dipolar signals corresponding to the distance distributions are shown in inserts.

In Figs. 12 and 13, we compare these SF-SVD results with those obtained from TIKR approaches (LongDistances and DeerLab), model fitting (DD Gaussian), and deep neural network (DEERNet). Fig. 12a shows the three monomer SF-SVD results from Fig. 11a for comparison with the results obtained from the other four methods (c.f. Fig.12 rows 2-5). Fig. 13a shows the three fibril SF-SVD results from Fig. 11b for comparison with the results obtained from the other four methods (c.f. Fig.13 rows 2-5). Starting with the monomer *P*(*r*) results, all five methods give similar distance distribution range and mean distance. The differences between the five methods are like that of the PEG results discussed in the previous section. The LongDistances results approximate those of SF-SVD but show less detail (e.g. Fig. 12 row 1 vs Fig. 12 row 2) and much greater uncertainty, which is due to the inherent difference in the way uncertainty is calculated. The DD Gaussian results (c.f. Fig. 12 row 3) captures subpopulation in the *P*(*r*), but also cannot resolve all the details because it tends to group small populations into a few gaussians. The DeerLab results captures more details than that of the LongDistance and DD Gaussian (c.f. Fig. 12 row 4); however, there are more spurious peaks in the broader distance distribution, specifically for the 291-322 distance because the regularization parameter automatically chosen can be suboptimal for some regions in the *P*(*r*). The DEERNet results give the smoothest *P*(*r*), with a small sharp peak around 4 nm for 300-313 monomer (c.f. Fig. 12b5). Because tau is an IDP and can have small features that are not present in polymers like PEG, methods that gives more detailed *P*(*r*) may be more optimal if extracting all distance subpopulations is the goal.

**Figure 12.**
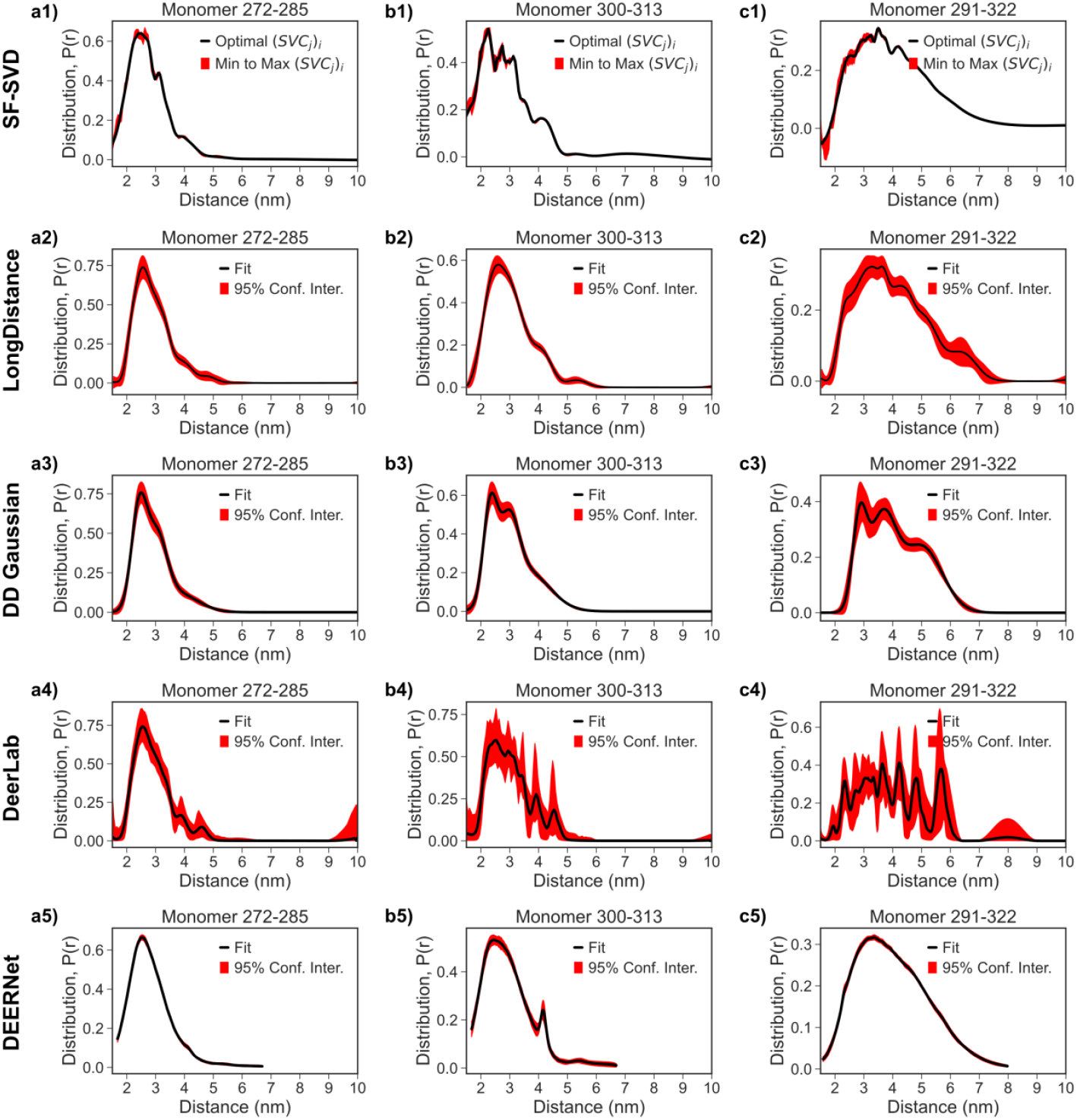
Comparison of Distance Distribution *P*(*r*) obtained from tau monomer DEER processed with the SF-SVD, LongDistance, DeerLab, DEERNet, and DD Gaussian methods. Column 1 **(a1-a5)** represents the tau monomer spin labeled at 272 and 285 amino acid sites. Column 2 **(b1-b5)** represents the tau monomer spin labeled at 300 and 313 amino acid sites. Column 3 **(c1-c5)** represents the tau monomer spin labeled at 291 and 322 amino acid sites. Row 1 **(a1, b1, c1)** represents *P*(*r*) from the SF-SVD method. Row 2 **(a2, b2, c2)** represents *P*(*r*) from the LongDistance method. Row 3 **(a3, b3, c3)** represents *P*(*r*) from the DD Gaussian method. Row 4 **(a4, b4, c4)** represents *P*(*r*) from the DeerLab method. Row 5 **(a5, b5, c5)** represents *P*(*r*) from the DEERNet method.

**Figure 13.**
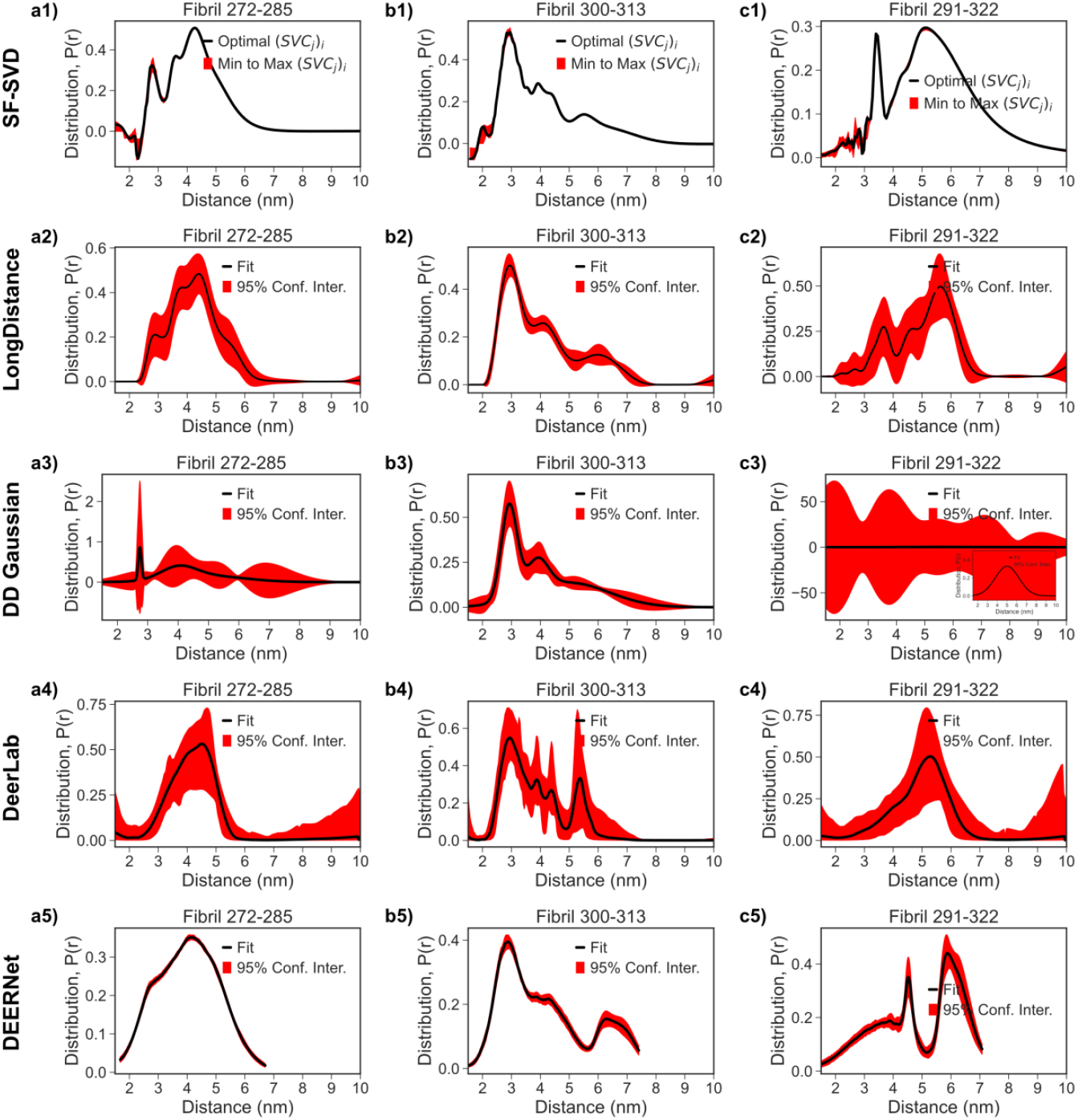
Comparison of Distance Distribution *P*(*r*) obtained from tau fibril DEER processed with the SF-SVD, LongDistance, DeerLab, DEERNet, and DD Gaussian methods. Column 1 **(a1-a5)** represents the tau fibril spin labeled at 272 and 285 amino acid sites. Column 2 **(b1-b5)** represents the tau fibril spin labeled at 300 and 313 amino acid sites. Column 3 **(c1-c5)** represents the tau fibril spin labeled at 291 and 322 amino acid sites. Row 1 **(a1, b1, c1)** represents *P*(*r*) from the SF-SVD method. Row 2 **(a2, b2, c2)** represents *P*(*r*) from the LongDistance method. Row 3 **(a3, b3, c3)** represents *P*(*r*) from the DD Gaussian method. Row 4 **(a4, b4, c4)** represents *P*(*r*) from the DeerLab method. Row 5 **(a5, b5, c5)** represents *P*(*r*) from the DEERNet method.

Looking at the fibril *P*(*r*) results, there are large differences between the five methods in how effectively they resolved narrow and board populations simultaneously. The LongDistances results approximate those of SF-SVD but show less detail (e.g. Fig. 13 row 1 vs Fig. 13 row 2). Specifically, for the 272-285 and 291-322 distances, the clearly resolved narrow distance population centered at 2.7 (cf. Fig. 13a1) and 3.6 nm (cf. Fig. 13c1), respectively, according to SF-SVD in the tau187 fibril state is blurred in the *P*(*r*) obtained from LongDistances, and hence would not be considered. The DD Gaussian results (c.f. Fig. 13 row 3) captures narrow and broad subpopulation in the *P*(*r*), but also cannot resolve all the details because it tends to group small populations into a few gaussians. Specifically, it failed to resolve the sharp narrow peak at 3.5 nm for 291-322 distance (c.f. Fig. 13c3) seen in the SF-SVD result (c.f. Fig. 13c1). The DeerLab results does not capture the same detail in the three previous mentioned methods, particularly evident for the 272-285 and 291-322 fibril distance distribution (cf. Fig. 13a4 and 13c4). Furthermore, the P(r) results from DeerLab appear to display extra spurious peaks in the *P*(*r*) for the 300-313 fibril distances (cf. Fig. 13b4). The discrepancies between results from the DeerLab result and the first three methods mean the one-step fitting approach is not suitable for our fibril data, which requires very careful background correction and careful choices of regularization parameter to resolve narrow peaks amidst broad peaks. However, with the same 2-step background correction approach and similar regularization parameter, the DeerLab result will be comparable to that of LongDistance but less resolved than that of SF-SVD. The DEERNet results are the least comparable to the other four methods, particularly evident for the 272-285 and 291-322 fibril distance distribution (cf. Fig. 13a5 and 13c5), where the subpopulations are over broadened and have different mean distances. This discrepancy means that P(r) with narrow and broad populations may not be the training dataset and DEERNet is currently not suitable for our tau fibril data. Base on this comparison, SF-SVD along with WavPDS is the most suitable for our tau fibril data.

## 4. CONCLUSION AND FUTURE WORK

We have shown the potential of DEER to study IDPs by applying WavPDS and SF-SVD. WavPDS can be exploited to collect the dipolar signals at longer evolution times, and so to yield more accurate *P*(*r*)′*s*. Most important, SF-SVD can reconstruct complex distance distributions associated with IDPs in a transparent manner and with no adjustable parameters, and hence in a user independent fashion. By offering a novel pseudo inversion approach of time-domain PDS data, in which the singular value cutoff can be selected separately for each distance, *r*, the SF-SVD method can determine and differentiate distance distributions containing distinct populations with multiple distances with simultaneous broad and narrow features, vs. a single broad distribution representing a dynamic ensemble.

Our future work will focus on other challenges in processing IDP data that includes identifying dimensionality for the spatial distribution of spin labels to achieve accurate baseline correction, improved baseline correction of dipolar signal that considers experimental artifacts, non-uniform sampling for increasing evolution time, applying denoising to raw, scan by scan, echo signals to further reduce the minimum SNR for WavPDS, quantifying the contribution of individual dipolar signal data at each distance distribution to identify effects of artifacts, and incorporating 2D SF-SVD methods to capture the structural evolution of IDPs into fibrils or other higher order assemblies.

## Acknowledgements

This research was supported by Cornell Startup Funds (M.S.), NIH grants P41GM103521 (J.H.F.), 1R24GM146107 (J.H.F.). S.H. and K.T. were supported by the National Institutes of Health under Grant R35GM136411 for dipolar EPR method developments. The study of key tau187 disease mutations was supported by the NIH grant R01AG05605 (S.H.).

