## Supplemental Information for "Localized Reconstruction of Multimodal Distance Distribution from DEER Data of Biopolymers"

| Table of Contents | Pages |
|---|---|
| **S.I.1** Data Processing Workflow. | 3-5 |
| **Table S1:** Comparison of regularization weights of different regularization criterion from DeerLab. | 6 |
| **Figure S1:** Comparison of distance distribution reconstructed from dipolar signal by Tikhonov regularization with different regularization parameters and that by SF-SVD using denoised dipolar signal. | 7 |
| **Figure S2:** PEG data processed with SF-SVD. | 8 |
| **Figure S3:** Tau monomer data processed with SF-SVD. | 9 |
| **Figure S4:** Tau fibril data processed with SF-SVD. | 10 |
| **Figure S5:** PEG data processed with LongDistance. | 11 |
| **Figure S6:** Tau monomer data processed with LongDistance. | 12 |
| **Figure S7:** Tau Fibril data processed with LongDistance. | 13 |
| **Figure S8:** PEG data processed with DeerLab. | 14 |
| **Figure S9:** Tau monomer data processed with DeerLab. | 15 |
| **Figure S10:** Tau fibril data processed with DeerLab. | 16 |
| **Figure S11:** PEG data processed with DD using Gaussian Model. | 17 |
| **Figure S12:** Tau Monomer processed with DD using Gaussian Model. | 18 |
| **Figure S13:** Tau fibril processed with DD using Gaussian Model. | 19 |
| **Figure S14:** PEG processed with DEERNet. | 20 |
| **Figure S15:** Tau monomer processed with DEERNet. | 21 |
| **Figure S16:** Tau fibril processed with DEERNet. | 22 |
| **References** | 23 |

### S.I.1 Data Processing Workflow

This section contains the detailed procedure and program link for the data processing present in this paper.

#### S.I.1.1 Data Denoising

Experimental DEER data, $V(t)$, was denoised using WavPDS (Wavelet-based Denoising for Pulsed Dipolar Signal). The data file was loaded into WavPDS as an ASCII text file with the time increment in the first column and signal amplitude in the second column. Wavelet family db and wavelet name db6 was chosen for all the samples in this paper. For each decomposition level, a threshold was chosen by picking the index number separating the low and high SNR region in the 'Detail component' plot. Threshold was chosen for each additional decomposition level, up to 7, until the data is properly denoised. The WavPDS program used is in the Denoising ESR Signals Vis Wavelets webpage hosted by the Cornell Center for Advanced Computing (CAC).[1]

#### S.I.1.2 SF-SVD

Denoised $V(t)$ is loaded into SVDReconstruction as an ASCII text file with the time increment in the first column and signal amplitude in the second column. Background correction was done according to Section 2.4 of the main text. Input signal was trimmed at time zero and the distance distribution was computed from 10 to 100 angstrom. Minimum and Maximum SVC number was chosen at the beginning and the end of the flat region of the linear scale Picard plot. Optimal SVC number was chosen between the minimum and maximum SVC number for the smoothest $P(r)$. For some samples, the ends of the signal were trimmed due to presence of artifacts from the signal itself or background correction. The $P(r)$ of some samples (i.e. tau fiber) contained multiple features; therefore, the $P(r)$ was divided into regions using the Breakpoints function and different SVC min and max was chosen for each region to represent uncertainty. The SVDReconstruction program used is in the Denoising ESR Signals Vis Wavelets webpage hosted by the Cornell Center for Advanced Computing (CAC).[2,3]

#### S.I.1.3 LongDistances

Raw $V(t)$ from Bruker Elexsys data file was loaded into Long Distances[4,5] for data analysis. For all samples but the tau fiber, 3D model: $B(t) = b_1 * e^{b_2 t}$, where $b_1$is the maximum of the signal and $b_2$ is the decay rate, was used to background correct the data. For the tau fiber, the dimensionality of the system was first determined by fitting the background only data using

VariableD model: $B(t) = b_1 * e^{[(b_2 t)^{\frac{b_3}{3}}]}$, where $b_3$ represents the dimensionality. Once determined, the dimensionality value was used in the VariableD model to perform background correction. Prior to background correction, the time traces were phase corrected and the end of the signal was eliminated if any artifact was present. The L curve was used as a criterion for determining the optimum smoothness parameter for the Model Free analysis by Tikhonov regularization for a smooth, but also a good fit to the $V(t)$. The uncertainty analysis was done using bootstrapping method with 100 samples. The distance distribution fits are presented with ±2 standard deviations (equivalent to 95% confidence interval).

**S.I.1.4 DeerLab**

Raw $V(t)$ from Bruker Elexsys data file was analyzed using DeerLab[6,7] software package for Python. The time traces were phase corrected and truncated to remove possible "2+1"-artifact. One-step analysis was done using the DeerLab fit function with the following models: ex-4deer model with t1, t2, and pulselength set to experiment parameters, bg-strexp model with the stretch parameter set to ~1.5/3 for tau fibril data (base on previous measurements of dimensions of singularly labeled samples) and 3/3 for all other data, and dipolarmodel using Tikhonov regularization. The regularization parameter was chosen automatically using the AIC criterion. The uncertainty analysis was done using bootstrapping method with 100 samples. The distance distribution fits are presented with 95% confidence interval.

**S.I.1.5 DD Gaussian**

Raw $V(t)$ from Bruker Elexsys data file was analyzed using DD software package for MATLAB. Using the DD MATLAB GUI, the time traces were phase corrected and the end of the signal was eliminated if any artifact was present. The background correction was performed the same way as the LongDistance method. The background corrected data was fitted with various number of gaussians, where the fit with the lowest reduced chi square was chosen. The uncertainty analysis was estimated from the variance-covariance matrix method. The distance distribution fits are presented with 95% confidence interval.

**S.I.1.6 DEERNet**

Raw $V(t)$ from Bruker Elexsys data file was analyzed using DEERNet[8–10] software package for MATLAB. Using the “elexsys2deernet” loading function, the time traces were automatically phase corrected and cropped to the echo maximum. For all PEG and tau monomer data, the following constraint was used: “bg_dim_range= [3.0,3.5]”. For all tau fibril data, the following constraint was used: “bg_dim_range= [2.0,3.5]”.  Using the “deernet” function, the data was automatically analyzed, and the function presents the distance distribution fit with 95% confidence interval.

| | **2.8 nm Ruler** | | **4.1 nm Ruler** | |
|---|---|---|---|---|
| | *Raw* | *Denoised* | *Raw* | *Denoised* |
| **AIC** | 0.027 | 0.023 | 0.456 | 0.339 |
| **AICC** | 0.023 | 0.027 | 0.479 | 0.341 |
| **BIC** | 0.045 | 0.048 | 0.762 | 0.522 |
| **CV** | 0.028 | 0.048 | 0.518 | 0.499 |
| **EE** | 0.017 | 0.0001 | 0.291 | 0.174 |
| **GCV** | 0.023 | 0.025 | 0.469 | 0.341 |
| **GML** | 0.030 | 0.023 | 0.800 | 0.348 |
| **LC** | 39810 | 0.794 | 100000 | 7943 |
| **LR** | 630 | 50 | 630 | 7.943 |
| **MCL** | 5942 | 49493 | 22177 | 98591 |
| **NCP** | 0.0001 | 0.0001 | 0.0003 | 0.148 |
| **RGVC** | 0.023 | 0.025 | 0.481 | 0.341 |
| **RM** | 15082 | 49569 | 67538 | 98976 |
| **SRGCV** | 11.69 | 5.49 | 25.53 | 3.802 |
| **L-Curve** | 2.83 | 1.16 | 9.28 | 3.81 |

**Table S1:** Comparison of regularization weights of different regularization criterion from DeerLab. Both the raw and denoised $V(t)$ of 2.8 nm Ruler and 4.1 nm Ruler was analyzed using DeerLab "dl.fit" function with the list of criterions from AIC to L-curve. The "dl.fit" function automatically optimizes for the regularization weight for each criterion and the values are reported.

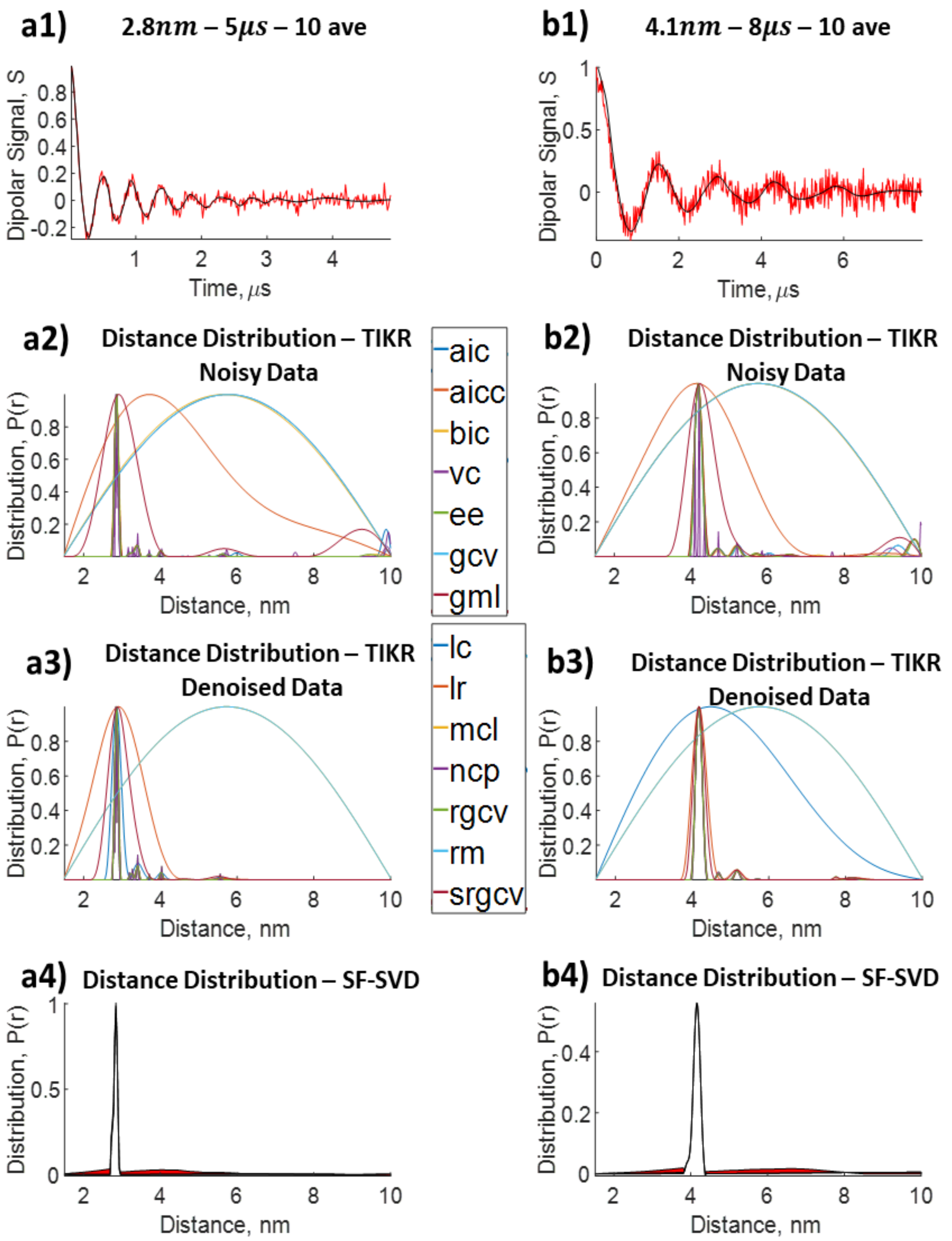


**Figure S1:** Comparison of distance distribution reconstructed from dipolar signal by Tikhonov regularization with different regularization parameters and that by SF-SVD using denoised dipolar signal. **a1)** 2.8$nm$ ruler with dipolar signal collected at 5$\mu s$ evolution time at 10 averages; **a2)** $P(r)$ from Tikhonov regularization for 2.8$nm$ ruler; **a3)** $P(r)$ from SF-TSVD for 2.8$nm$ ruler; **b1)** 4.1$nm$ ruler with dipolar signal collected at 5$\mu s$ evolution time at 10 averages; **b2)** $P(r)$ from Tikhonov regularization for 4.1$nm$ ruler; **b3)** $P(r)$ from SF-TSVD for 4.1$nm$ ruler.

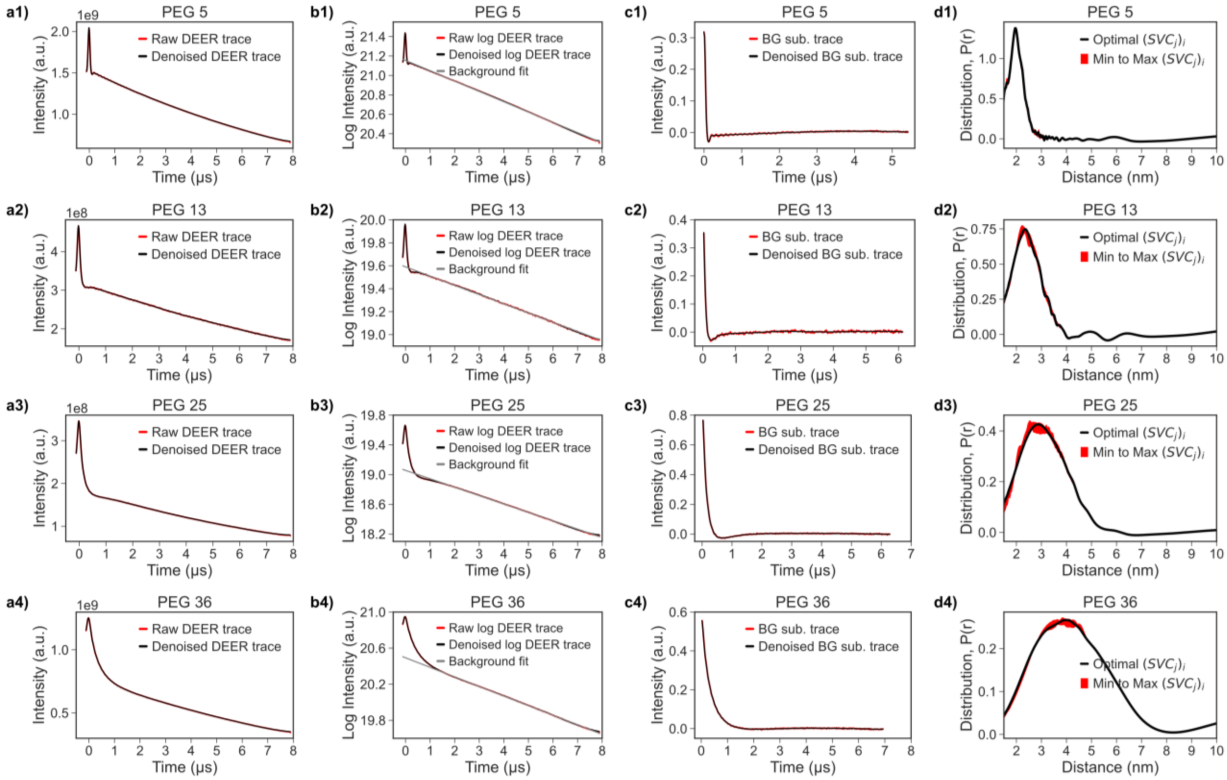


**Figure S2:** PEG data processed with SF-SVD. Row 1 (a1, b1, c1, d1) represents end-to-end labeled PEG 5. Row 2 (a2, b2, c2, d2) represents end-to-end labeled PEG 13. Row 3 (a3, b3, c3, d3) represents end-to-end labeled PEG 25. Row 4 (a4, b4, c4, d4) represents end-to-end labeled PEG 36. Column 1 (a1-a4) represents the raw DEER trace and the denoised DEER trace. Column 2 (b1-b4) represents the log of raw DEER trace, denoised DEER trace, and the background fit. Column 3 (c1-c4) represents the antilog of the raw and denoised DEER trace after background subtraction in log domain. Column 4 (d1-d4) represents distance distribution from the SF-SVD method, with the uncertainty defined as minimum and maximum SVC number.

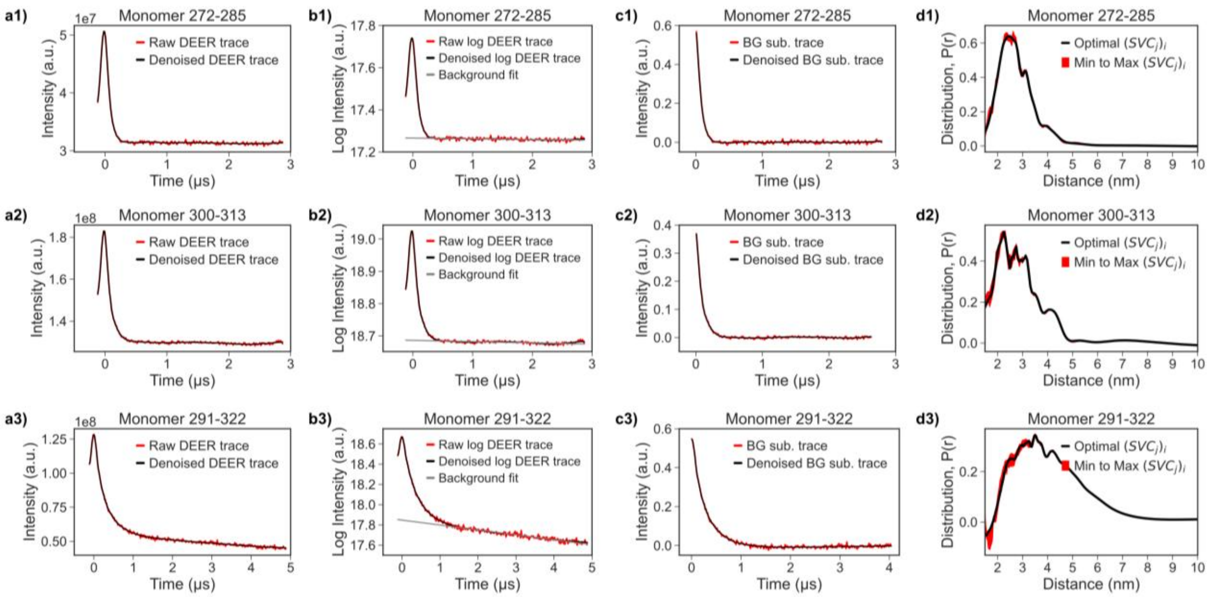


**Figure S3:** Tau monomer data processed with SF-SVD. Row 1 (a1, b1, c1, d1) represents the tau monomer spin labeled at 272 and 285 amino acid sites. Row 2 (a2, b2, c2, d2) represents the tau monomer spin labeled at 300 and 313 amino acid sites. Row 3 (a3, b3, c3, d3) represents the tau monomer spin labeled at 291 and 322 amino acid sites. Column 1 (a1-a3) represents the raw DEER trace and the denoised DEER trace. Column 2 (b1-b3) represents the log of raw DEER trace, denoised DEER trace, and the background fit. Column 3 (c1-c3) represents the antilog of the raw and denoised DEER trace after background subtraction in log domain. Column 4 (d1-d3) represents distance distribution from the SF-SVD method, with the uncertainty defined as minimum and maximum SVC number.

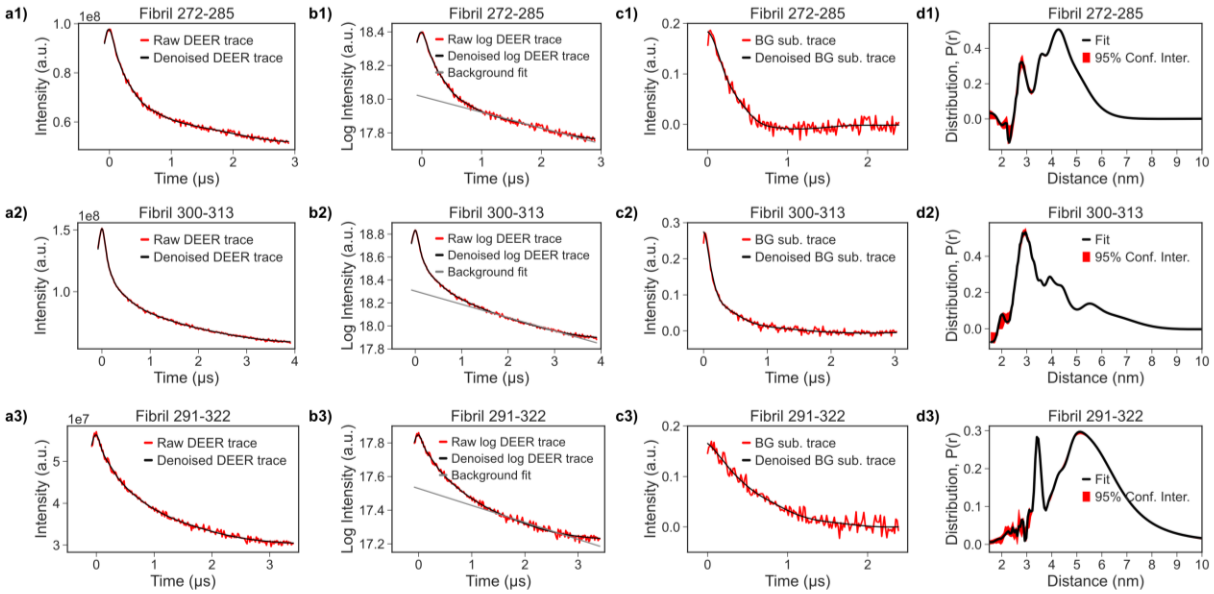


**Figure S4:** Tau fibril data processed with SF-SVD. Row 1 (a1, b1, c1, d1) represents the tau fibril spin labeled at 272 and 285 amino acid sites. Row 2 (a2, b2, c2, d2) represents the tau fibril spin labeled at 300 and 313 amino acid sites. Row 3 (a3, b3, c3, d3) represents the tau fibril spin labeled at 291 and 322 amino acid sites. Column 1 (a1-a3) represents the raw DEER trace and the denoised DEER trace. Column 2 (b1-b3) represents the log of raw DEER trace, denoised DEER trace, and the background fit. Column 3 (c1-c3) represents the antilog of the raw and denoised DEER trace after background subtraction in log domain. Column 4 (d1-d3) represents distance distribution from the SF-SVD method, with the uncertainty defined as minimum and maximum SVC number.

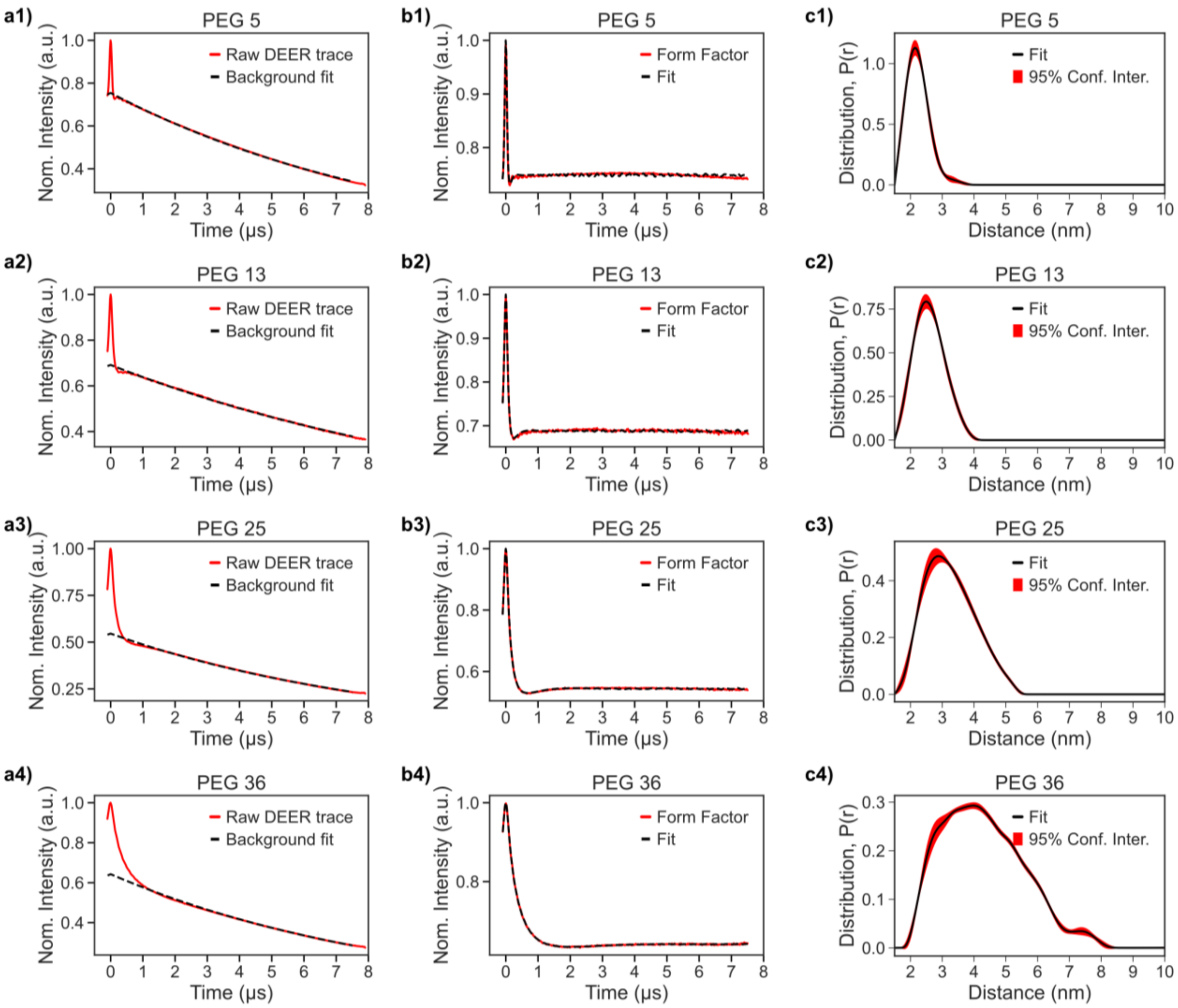


**Figure S5:** PEG data processed with LongDistance. PEG data processed with DD using Gaussian Model. Row 1 (a1, b1, c1) represents end-to-end labeled PEG 5. Row 2 (a2, b2, c2) represents end-to-end labeled PEG 13. Row 3 (a3, b3, c3) represents end-to-end labeled PEG 25. Row 4 (a4, b4, c4) represents end-to-end labeled PEG 36. Column 1 (a1-a4) represents the raw DEER trace and its background fit. Column 2 (b1-b4) represents the form factor (trimmed DEER trace after background division) and its fit. Column 3 (c1-c4) represents the distance distribution with 95% confidence interval.

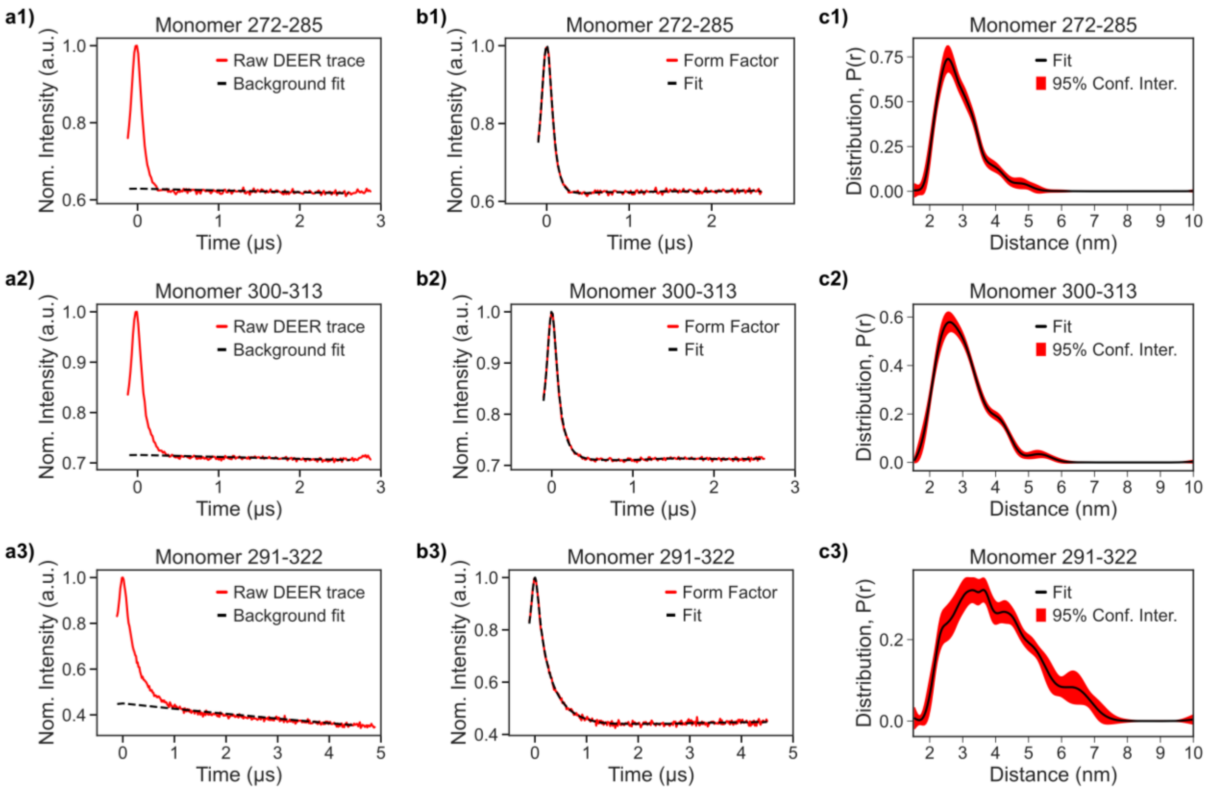


**Figure S6:** Tau monomer data processed with LongDistance. Row 1 (a1, b1, c1) represents the tau monomer spin labeled at 272 and 285 amino acid sites. Row 2 (a2, b2, c2) represents the tau monomer spin labeled at 300 and 313 amino acid sites. Row 3 (a3, b3, c3) represents the tau monomer spin labeled at 291 and 322 amino acid sites. Column 1 (a1-a3) represents the raw DEER trace and its background fit. Column 2 (b1-b3) represents the form factor (trimmed DEER trace after background division) and its fit. Column 3 (c1-c3) represents the distance distribution with 95% confidence interval.

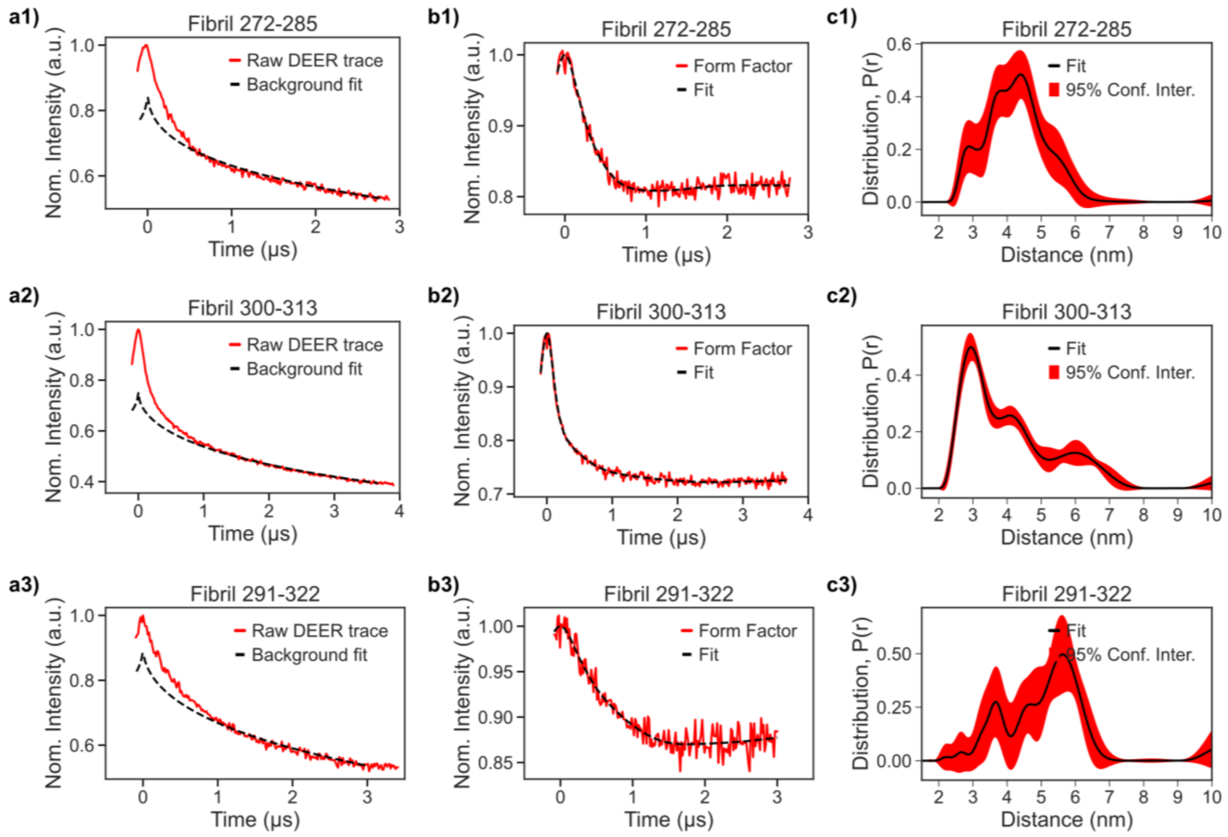


**Figure S7:** Tau Fibril data processed with LongDistance. Row 1 (a1, b1, c1) represents the tau fibril spin labeled at 272 and 285 amino acid sites. Row 2 (a2, b2, c2) represents the tau fibril spin labeled at 300 and 313 amino acid sites. Row 3 (a3, b3, c3) represents the tau fibril spin labeled at 291 and 322 amino acid sites. Column 1 (a1-a3) represents the raw DEER trace and its background fit. Column 2 (b1-b3) represents the form factor (trimmed DEER trace after background division) and its fit. Column 3 (c1-c3) represents the distance distribution with 95% confidence interval.

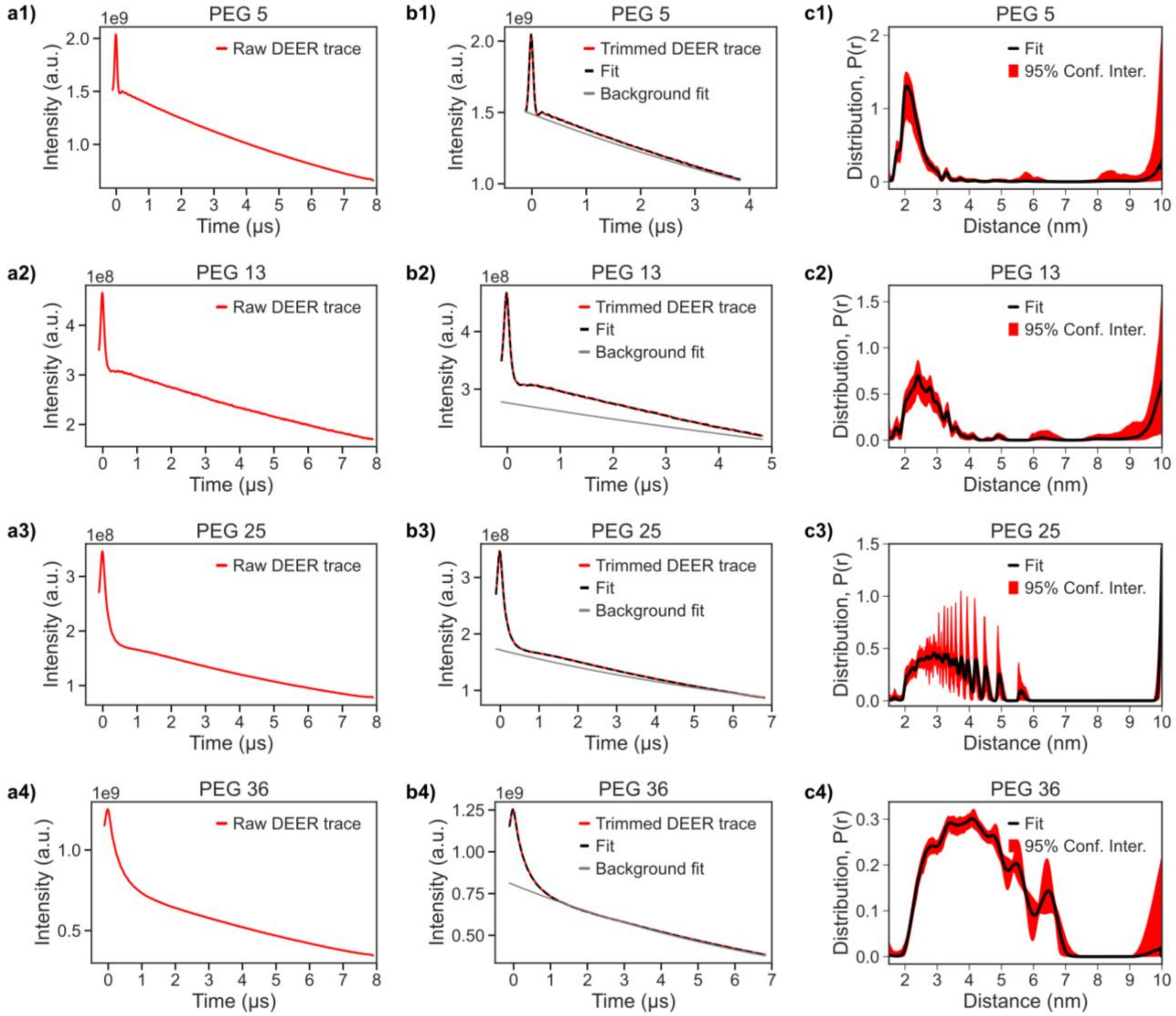


**Figure S8:** PEG data processed with DeerLab. Row 1 (a1, b1, c1) represents end-to-end labeled PEG 5. Row 2 (a2, b2, c2) represents end-to-end labeled PEG 13. Row 3 (a3, b3, c3) represents end-to-end labeled PEG 25. Row 4 (a4, b4, c4) represents end-to-end labeled PEG 36. Column 1 (a1-a4) represents the raw DEER trace. Column 2 (b1-b4) represents the trimmed DEER trace, background fit, and DEER trace fit. Column 3 (c1-c4) represents the distance distribution with 95% confidence interval.

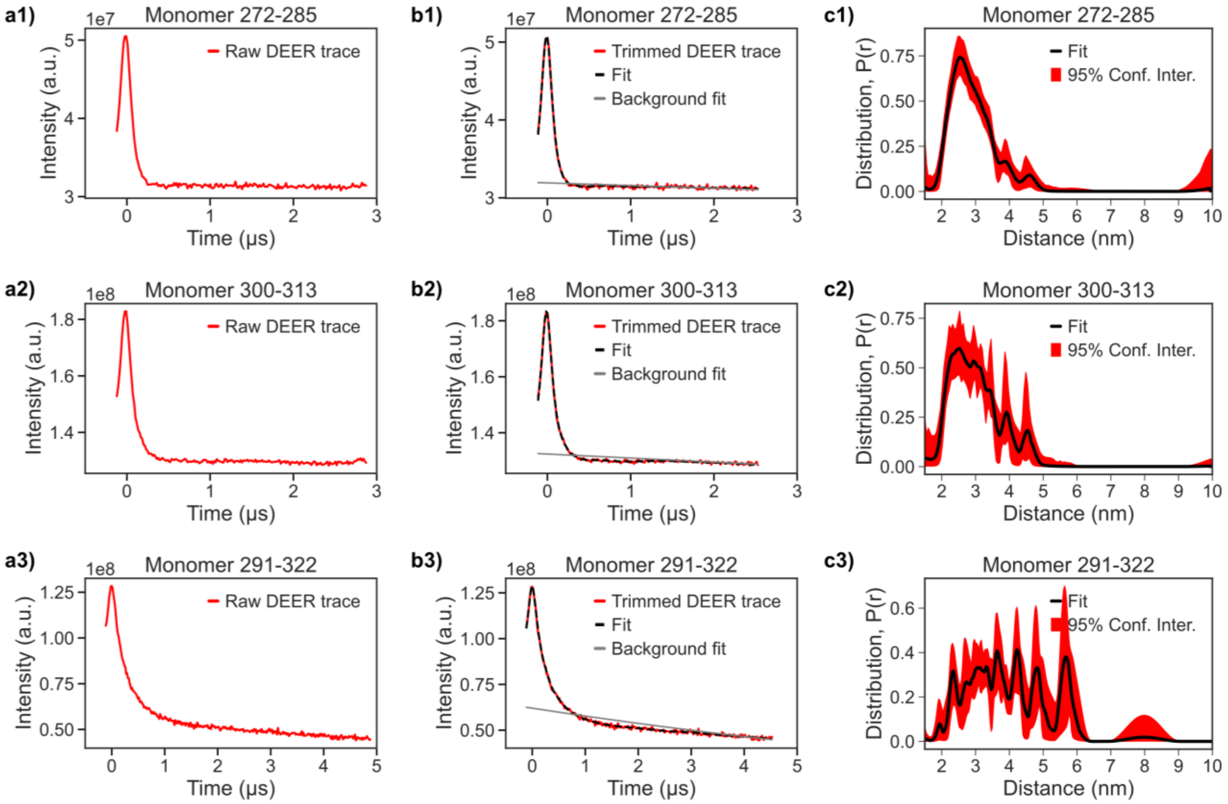


**Figure S9:** Tau monomer data processed with DeerLab. Row 1 (a1, b1, c1) represents the tau monomer spin labeled at 272 and 285 amino acid sites. Row 2 (a2, b2, c2) represents the tau monomer spin labeled at 300 and 313 amino acid sites. Row 3 (a3, b3, c3) represents the tau monomer spin labeled at 291 and 322 amino acid sites. Column 1 (a1-a4) represents the raw DEER trace. Column 2 (b1-b4) represents the trimmed DEER trace, background fit, and DEER trace fit. Column 3 (c1-c4) represents the distance distribution with 95% confidence interval.

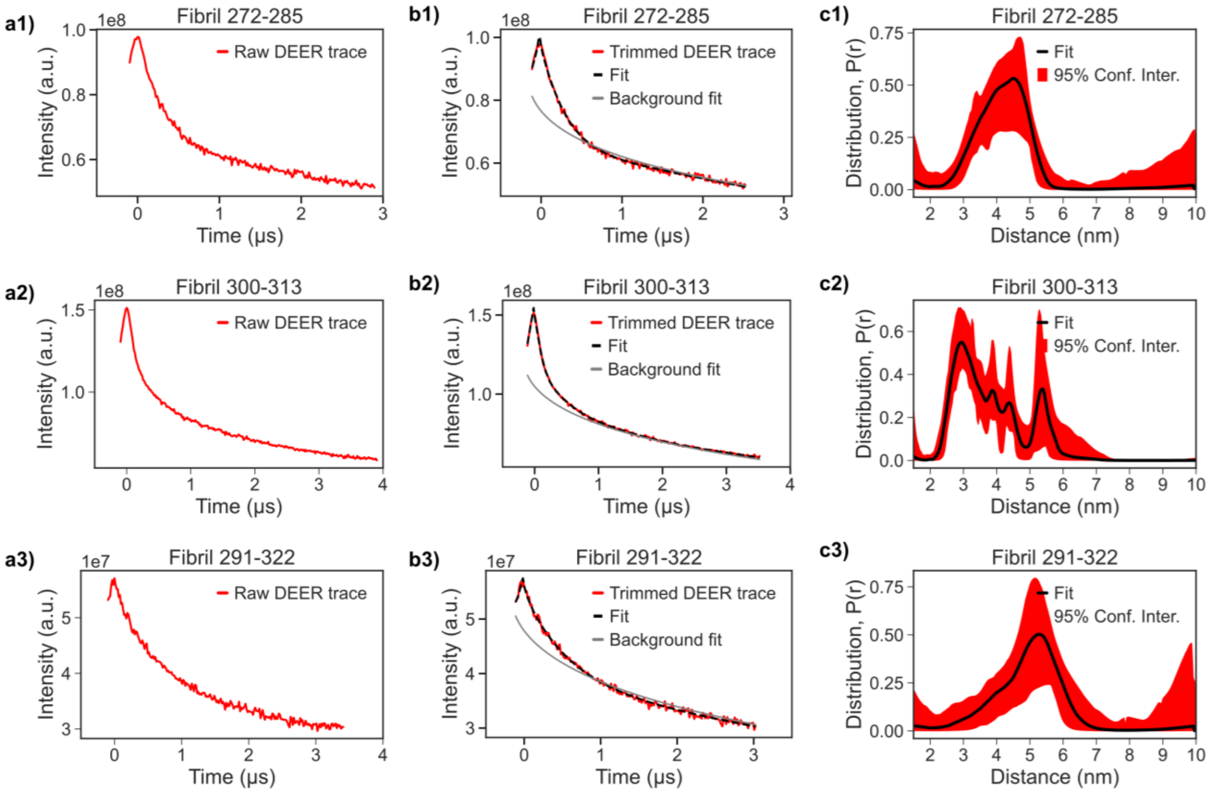


**Figure S10:** Tau fibril data processed with DeerLab. Row 1 (a1, b1, c1) represents the tau fibril spin labeled at 272 and 285 amino acid sites. Row 2 (a2, b2, c2) represents the tau fibril spin labeled at 300 and 313 amino acid sites. Row 3 (a3, b3, c3) represents the tau fibril spin labeled at 291 and 322 amino acid sites. Column 1 (a1-a4) represents the raw DEER trace. Column 2 (b1-b4) represents the trimmed DEER trace, background fit, and DEER trace fit. Column 3 (c1-c4) represents the distance distribution with 95% confidence interval.

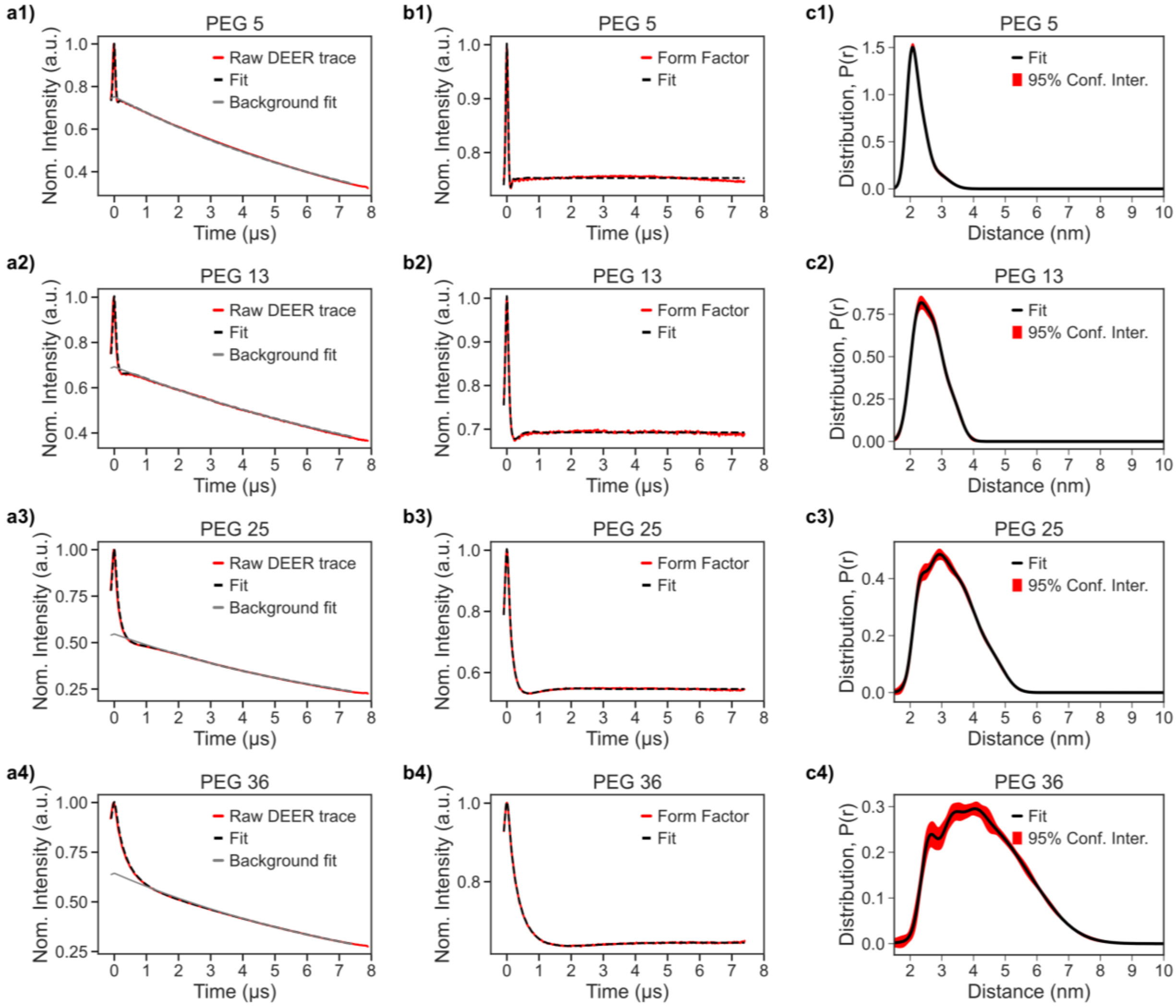


**Figure S11:** PEG data processed with DD using Gaussian Model. Row 1 (a1, b1, c1) represents end-to-end labeled PEG 5. Row 2 (a2, b2, c2) represents end-to-end labeled PEG 13. Row 3 (a3, b3, c3) represents end-to-end labeled PEG 25. Row 4 (a4, b4, c4) represents end-to-end labeled PEG 36. Column 1 (a1-a4) represents the raw DEER trace, fit of the trimmed DEER trace, and background fit of the trimmed DEER trace. Column 2 (b1-b4) represents the form factor (trimmed DEER trace after background division) and its fit. Column 3 (c1-c4) represents the distance distribution with 95% confidence interval.

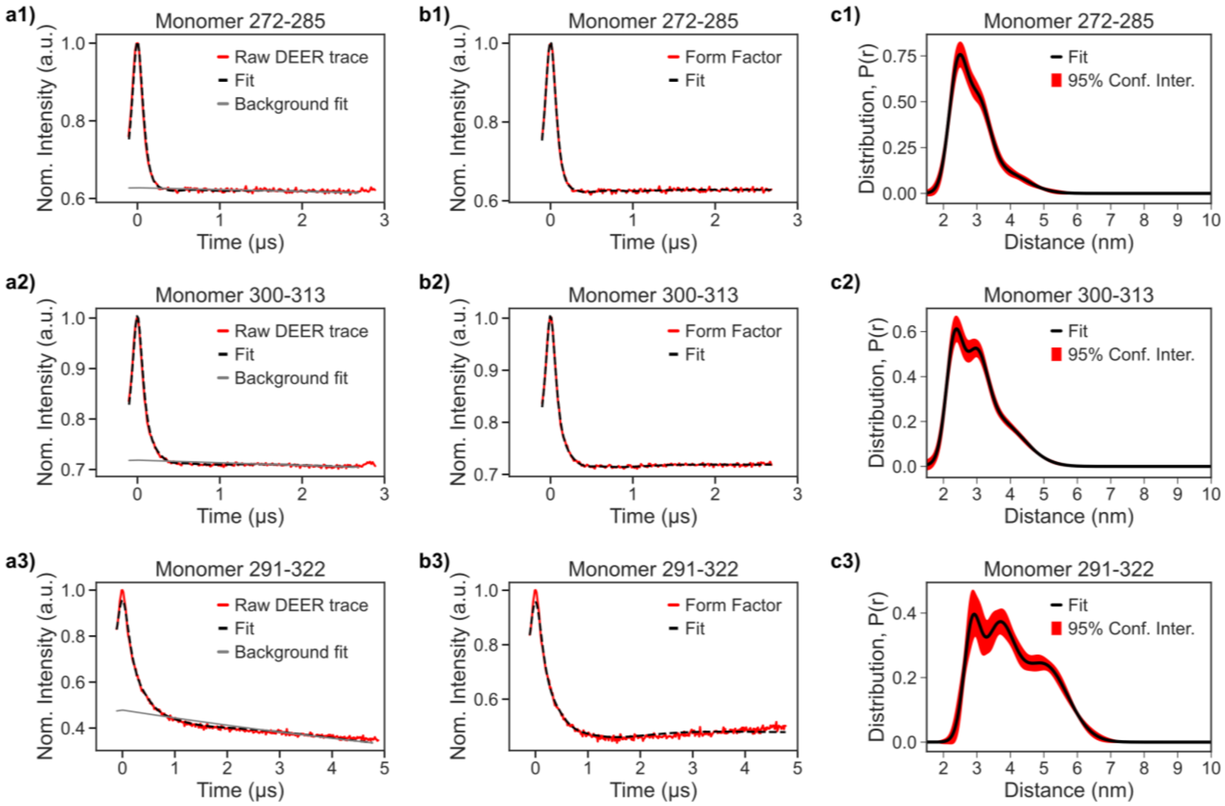


**Figure S12:** Tau Monomer processed with DD using Gaussian Model. Row 1 (a1, b1, c1) represents the tau monomer spin labeled at 272 and 285 amino acid sites. Row 2 (a2, b2, c2) represents the tau monomer spin labeled at 300 and 313 amino acid sites. Row 3 (a3, b3, c3) represents the tau monomer spin labeled at 291 and 322 amino acid sites. Column 1 (a1-a3) represents the raw DEER trace, fit of the trimmed DEER trace, and background fit of the trimmed DEER trace. Column 2 (b1-b3) represents the form factor (trimmed DEER trace after background division) and its fit. Column 3 (c1-c3) represents the distance distribution with 95% confidence interval.

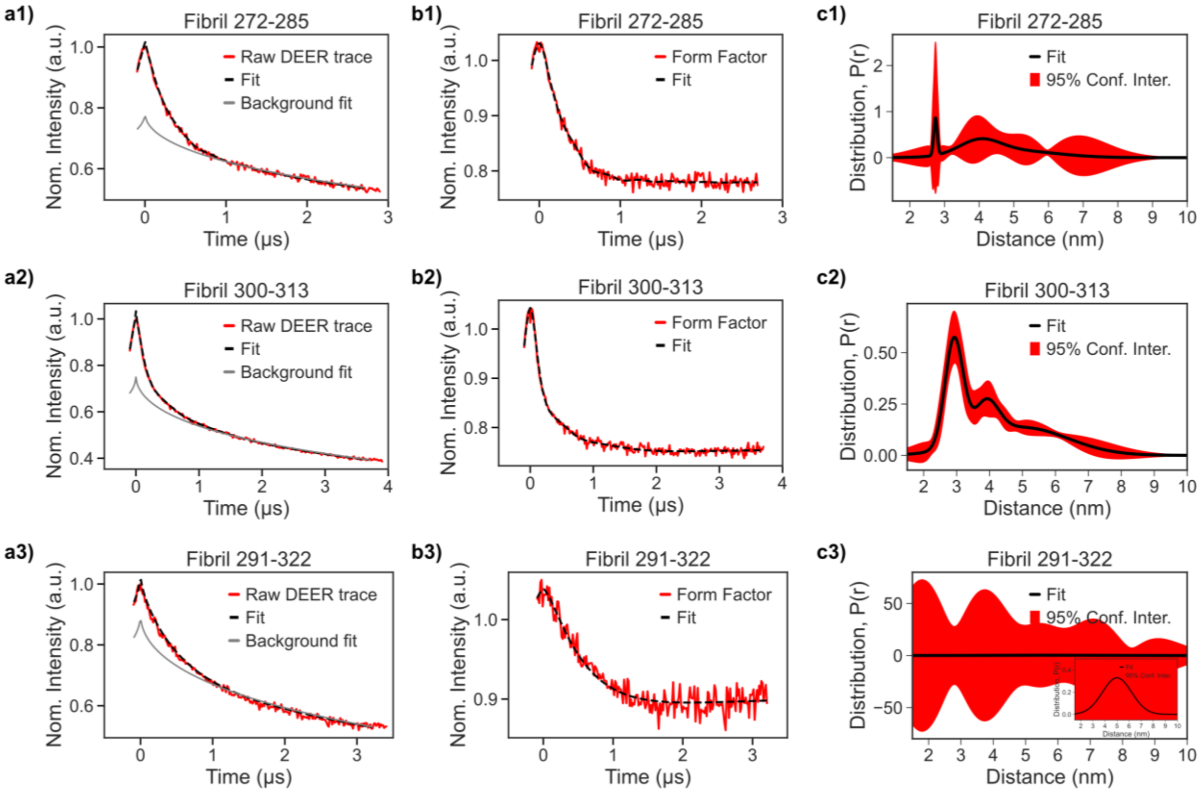


**Figure S13:** Tau fibril processed with DD using Gaussian Model. Row 1 (a1, b1, c1) represents the tau fibril spin labeled at 272 and 285 amino acid sites. Row 2 (a2, b2, c2) represents the tau fibril spin labeled at 300 and 313 amino acid sites. Row 3 (a3, b3, c3) represents the tau fibril spin labeled at 291 and 322 amino acid sites. Column 1 (a1-a3) represents the raw DEER trace, fit of the trimmed DEER trace, and background fit of the trimmed DEER trace. Column 2 (b1-b3) represents the form factor (trimmed DEER trace after background division) and its fit. Column 3 (c1-c3) represents the distance distribution with 95% confidence interval.

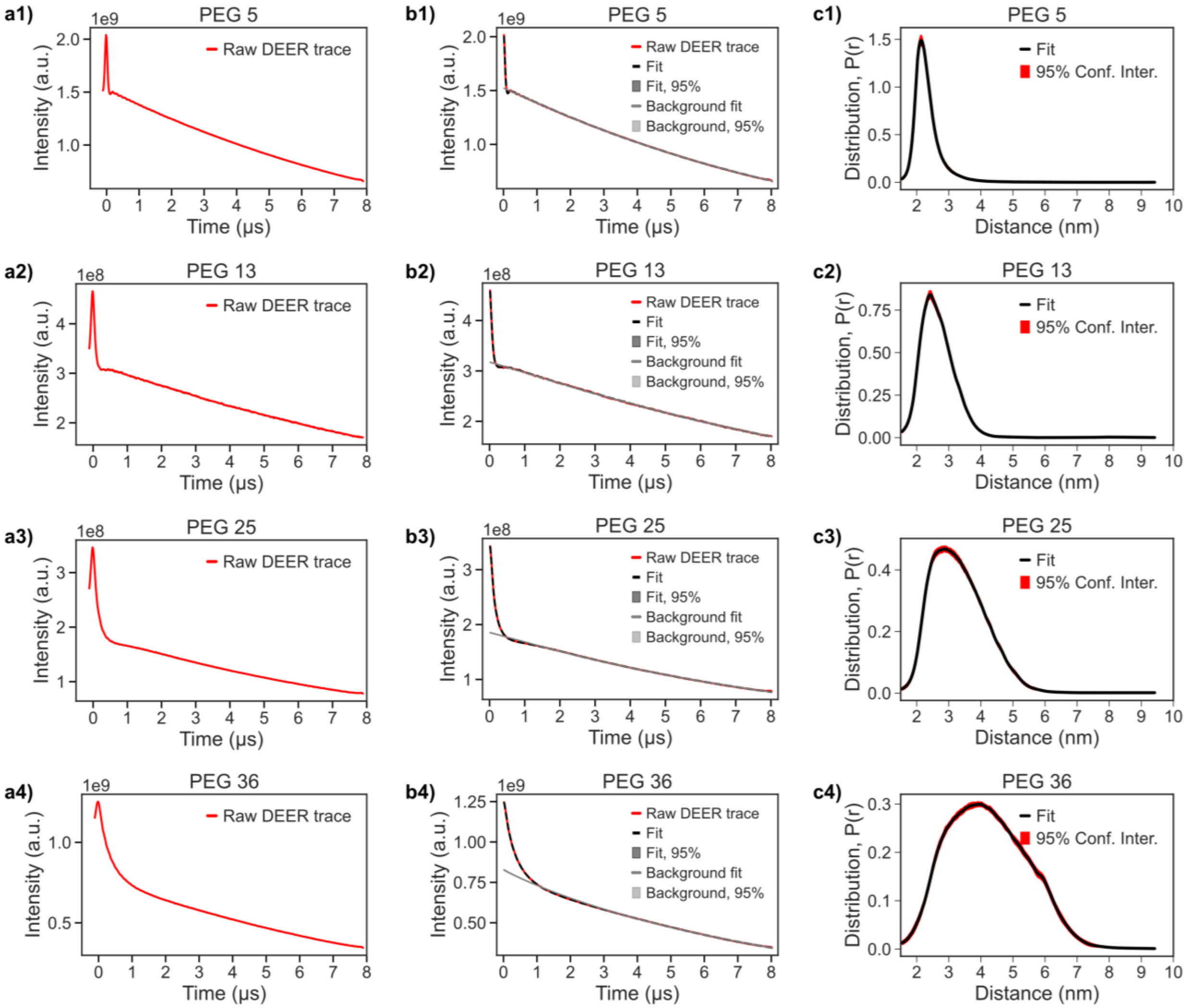


**Figure S14:** PEG processed with DEERNet. PEG data processed with DD using Gaussian Model. Row 1 (a1, b1, c1) represents end-to-end labeled PEG 5. Row 2 (a2, b2, c2) represents end-to-end labeled PEG 13. Row 3 (a3, b3, c3) represents end-to-end labeled PEG 25. Row 4 (a4, b4, c4) represents end-to-end labeled PEG 36. Column 1 (a1-a4) represents the raw DEER trace. Column 2 (b1-b4) represents the raw DEER trace, background fit, and DEER trace fit. Column 3 (c1-c4) represents the distance distribution with 95% confidence interval.

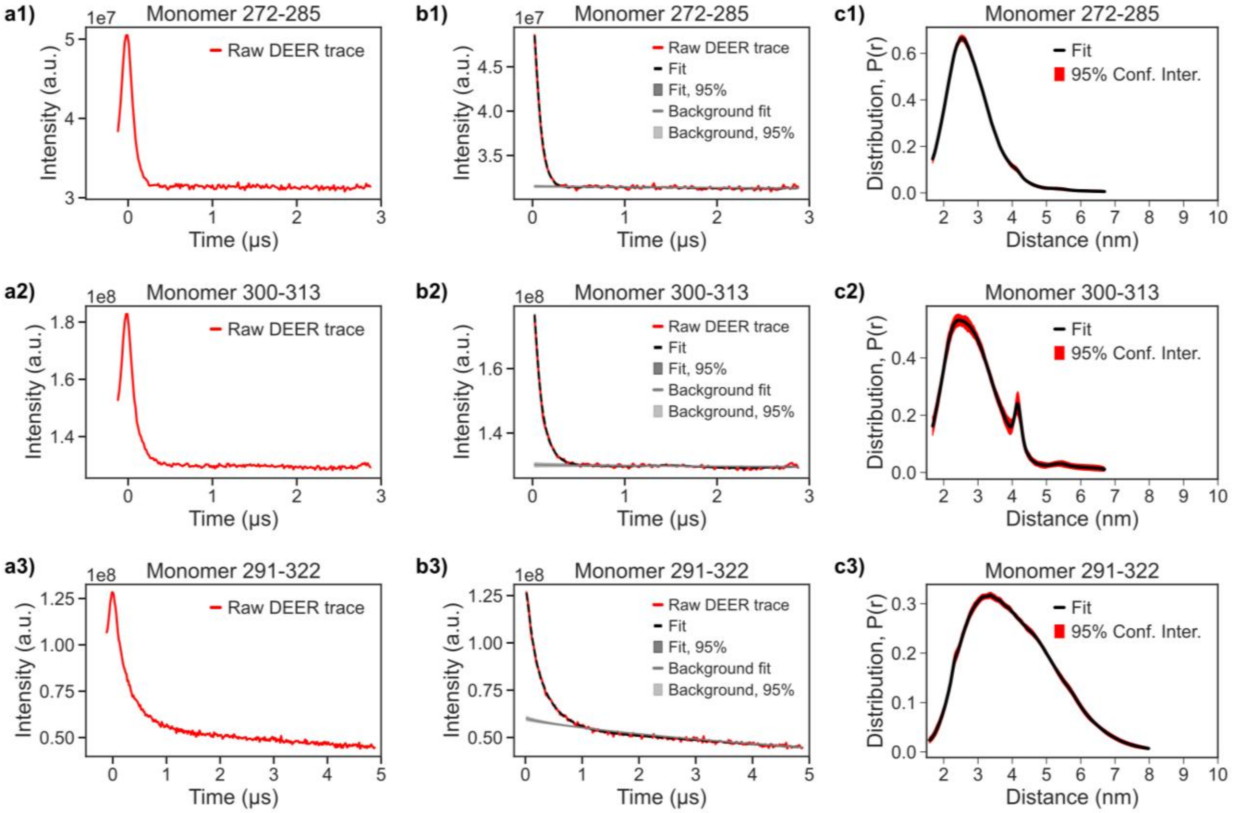


**Figure S15:** Tau monomer processed with DEERNet. Row 1 (a1, b1, c1) represents the tau monomer spin labeled at 272 and 285 amino acid sites. Row 2 (a2, b2, c2) represents the tau monomer spin labeled at 300 and 313 amino acid sites. Row 3 (a3, b3, c3) represents the tau monomer spin labeled at 291 and 322 amino acid sites. Column 1 (a1-a3) represents the raw DEER trace. Column 2 (b1-b3) represents the raw DEER trace, background fit, and DEER trace fit. Column 3 (c1-c3) represents the distance distribution with 95% confidence interval.

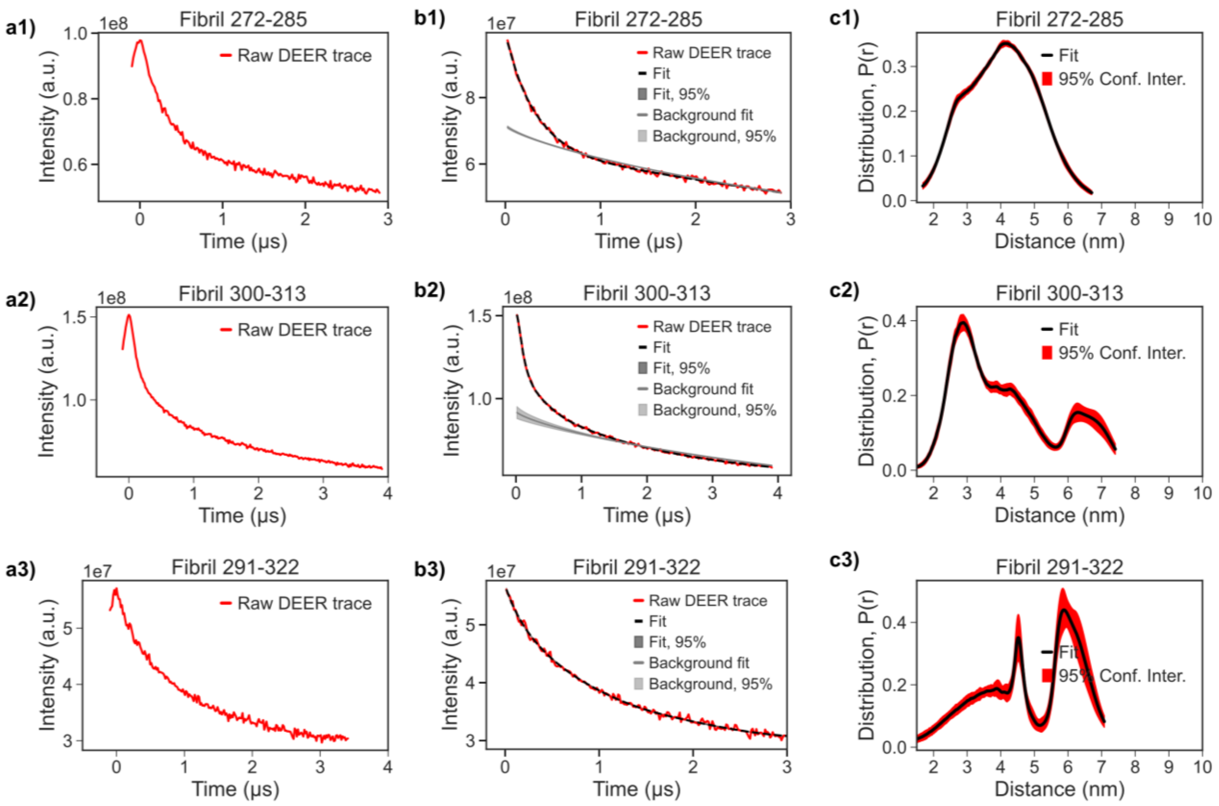


**Figure S16:** Tau fibril processed with DEERNet. Row 1 (a1, b1, c1) represents the tau fibril spin labeled at 272 and 285 amino acid sites. Row 2 (a2, b2, c2) represents the tau fibril spin labeled at 300 and 313 amino acid sites. Row 3 (a3, b3, c3) represents the tau fibril spin labeled at 291 and 322 amino acid sites. Column 1 (a1-a3) represents the raw DEER trace. Column 2 (b1-b3) represents the raw DEER trace, background fit, and DEER trace fit. Column 3 (c1-c3) represents the distance distribution with 95% confidence interval.
